# *De Novo* PacBio long-read and phased avian genome assemblies correct and add to genes important in neuroscience research

**DOI:** 10.1101/103911

**Authors:** Jonas Korlach, Gregory Gedman, Sarah B. Kingan, Chen-Shan Chin, Jason Howard, Lindsey Cantin, Erich D. Jarvis

## Abstract

Reference quality genomes are expected to provide a resource for studying gene structure and function. However, often genes of interest are not completely or accurately assembled, leading to unknown errors in analyses or additional cloning efforts for the correct sequences. A promising solution to this problem is long-read sequencing. Here we tested PacBio-based long-read sequencing and diploid assembly for potential improvements to the Sanger-based intermediate-read zebra finch reference and Illumina-based short-read Anna’s hummingbird reference, two vocal learning avian species widely studied in neuroscience and genomics. With DNA of the same individuals used to generate the reference genomes, we generated diploid assemblies with the FALCON-Unzip assembler, resulting in contigs with no gaps in the megabase range (N50s of 5.4 and 7.7 Mb, respectively), and representing 150-fold and 200-fold improvements over the current zebra finch and hummingbird references, respectively. These long-read assemblies corrected and resolved what we discovered to be misassemblies, including due to erroneous sequences flanking gaps, complex repeat structure errors in the references, base call errors in difficult to sequence regions, and inaccurate resolution of allelic differences between the two haplotypes. We analyzed protein-coding genes widely studied in neuroscience and specialized in vocal learning species, and found numerous assembly and sequence errors in the reference genes that the PacBio-based assemblies resolved completely, validated by single long genomic reads and transcriptome reads. These findings demonstrate, for the first time in non-human vocal learning species, the impact of higher quality, phased and gap-less assemblies for understanding gene structure and function.

## Introduction

Having an available genome of a species of interest provides a powerful resource to rapidly conduct investigations on genes of interest. For example, using the costly Sanger method to sequence genomes of the two most commonly studied bird species, the chicken (Hillier et al. 2004) and zebra finch (Warren et al. 2010), have impacted many studies. The zebra finch is also a vocal learning songbird, with the rare ability to imitate sounds as humans do for speech; comparative analyses of genes in its genome has allowed insights into the mechanisms and evolution of spoken-language in humans (Warren et al. 2010; Shi et al. 2013; Pfenning et al. 2014). Subsequently with the advent of more cost-effective next generation sequencing technologies using short reads, many new genomes were sequenced, with one large successful project conducted by the Avian Phylogenomics Consortium, generating genomes of 45 new bird species and several reptiles (Zhang et al. 2014b). The consortium was very successful in conducting comparative genomics and phylogenetics with populations of genes (Jarvis et al. 2014; Zhang et al. 2014c; Joseph and Buchanan 2015; Kraus and Wink 2015). However, when it was necessary to dig deeper to study individual genes, it was discovered that many were incompletely assembled or contain apparent misassemblies. For example, the *DRD4* dopamine receptor was missing in half of the assemblies, in part due to sequence complexity (Haug-Baltzell et al. 2015). The *EGR1* immediate early gene transcription factor, a commonly studied gene in neuroscience and in vocal learning species, was missing the promoter region in an apparent GC-rich region in every bird genome we examined. Another immediate early gene, *DUSP1*, with specialized vocalizing-driven gene expression in song nuclei of vocal learning species, has microsatellite sequences in the promoters of vocal learning species that are missing or misassembled, requiring single-molecule cloning to resolve the genes (Horita et al. 2012). Such errors create a great amount of effort to clone, sequence, and correct assemblies of individual genes of interest.

High-throughput, single-molecule, long-read sequencing shows promise to alleviate these problems (Bradnam et al. 2013; Roberts et al. 2013; Gordon et al. 2016). Here, we applied PacBio single-molecule long-read (1000-60,000 bp) sequencing and diploid assembly on two vocal learning species, the zebra finch previously assembled with Sanger-based intermediate reads (700-1000 bp), and Anna’s hummingbird previously assembled with Illumina-based short reads (100-150 bp). We found that the long-read diploid assemblies resulted in major improvements in genome completeness and contiguity, and completely resolved the problems in all of our genes of interest. This study is part of an effort to help evaluate standards for the G10K vertebrate (https://genome10k.soe.ucsc.edu) and the B10K bird (http://b10k.genomics.cn/index.html) genome consortiums.

## Results and Discussion

### The long-read assemblies show 150-fold to 200-fold increases in contiguity

High molecular weight DNA was isolated from muscle tissue of the same zebra finch male and Anna’s hummingbird female used to create the current reference genomes (Warren et al. 2010; Zhang et al. 2014c). The DNA was sheared, 35-40 kb libraries generated, size-selected for inserts >17 kb (**Suppl. Fig. 1**), and then SMRT sequencing performed on the PacBio RS II instrument to obtain ~96X coverage for the zebra finch (19 kb read length N50) and ~70X for the hummingbird (22 kb read length N50; **Suppl. Fig. 2**). The long reads were originally assembled into a merged haplotype with an early version of the FALCON assembler (Chin 2015), which we found unintentionally introduced indels for some nucleotides that differed between haplotypes (tested on the hummingbird; data not shown). We then re-assembled using FALCON v0.4.0 followed by the FALCON-Unzip module (Chin et al. 2016) to prevent indel formation and generate long-range phased haplotypes. Thus, the new assemblies, unlike the current reference assemblies, are phased diploid. This PacBio-based sequencing and assembly approach does not link contigs into gapped scaffolds; scaffolding requires additional approaches, which we will report on separately in a study comparing scaffolding technologies with these assemblies. The results presented here were found independent of scaffolding.

For the zebra finch, our long-read approach resulted in 1159 primary haplotype contigs with an estimated total genome size of 1.14 Gb (1.2 Gb expected; http://www.genomesize.com/results.php?page=1) and contig N50 of 5.81 Mb, representing a 108-fold reduction in the number of contigs and a 150-fold improvement in contiguity compared to the current Sanger-based reference (**Table 1A**). The diploid assembly process produced 2188 associated, or secondary, haplotype contigs (i.e. haplotigs) with an estimated length of 0.84 Gb (**Table 1A**), implying that about 75% of the genome contained sufficient heterozygosity to be phased into haplotypes by FALCON-Unzip. Since in FALCON-Unzip, the primary contigs are the longest path through the assembly string graph, the secondary haplotigs are by definition shorter and can be more numerous, resulting in lower contiguity for the haplotigs. Regions of the genome with very low heterozygosity remain as collapsed haplotypes in the primary contigs.

The PacBio long-read assembly for the hummingbird was of similar quality, with 1076 primary contigs generating a primary haploid genome size of 1.01 Gb (1.14 Gb expected; http://www.genomesize.com/results.php?page=1), and a contig N50 of 5.36 Mb, representing a 116-fold reduction in the number of contigs and a 201-fold improvement in contiguity over the current reference (**Table 1B**). The length of the assembled secondary haplotigs for the hummingbird was similar to that of the primary contig backbone (1.01 Gb; **Table 1B**) indicating that there was sufficient heterozygosity to phase most of the diploid genome into the two haplotypes.

**Table 1:** *De novo* genome assembly statistics comparing intermediate-read length and short-read length assemblies with the long-read assemblies. (A) Zebra finch intermediate-read length (Sanger-based, NCBI accession # GCF_000151805, version 3.2.4) compared to the long-read length PacBio-based assembly. (B) Anna’s hummingbird short-read length (Illumina-based, accession # GCF_000699085) compared to the long-read length PacBio-based assembly. Improvement is calculated between the 2^nd^ and 3^rd^ columns for the primary PacBio-based haplotype. The higher number of contigs in the secondary haplotype (5^th^ column) are a result of the arbitrary assignment of shorter haplotypes to the haplotig category.

| Species | Reference assembly | PacBio-based primary haplotype | Improvement | PacBio-based secondary haplotype |
| --- | --- | --- | --- | --- |
| **A. Zebra finch** | **Sanger-based** |  |  |  |
| Number of contigs | 124,806 | 1,159 | - 108 fold | 2,188 |
| Contig N50 | 38,639 bp | 5,807,022 bp | + 150 fold | 2,740,176 bp |
| Total size | 1,232,135,591 bp | 1,138,770,338 bp |  | 843,915,757 bp |
| **B. Hummingbird** | **Illumina-based** |  |  |  |
| Number of contigs | 124,820 | 1,076 | - 116 fold | 4,895 |
| Contig N50 | 26,738 bp | 5,366,327 bp | + 201 fold | 1,073,631 bp |
| Total size | 1,105,676,412 bp | 1,007,374,986 bp |  | 1,013,746,550 bp |

### The long-read assemblies have more complete conserved protein coding genes

To assess gene completeness, we analyzed 248 highly conserved eukaryotic genes from the CEGMA human set (Parra et al. 2007; Parra et al. 2009) in each of the assemblies. Both the PacBio-based zebra finch and hummingbird assemblies showed improved resolution of these genes, with a close to doubling (~71%) for the zebra finch and 26% increase for the hummingbird in the number of complete or near-complete (>95%) CEGMA genes assembled, compared to the references (**Fig. 1A**). Because updating the CEGMA gene sets was recently discontinued due to lack of continued funding and ease of use (http://www.acgt.me/blog/2015/5/18/goodbye-cegma-hello-busco), we also searched for a set of conserved, single-copy genes from the orthoDB9 (Zdobnov et al. 2017) gene set using the recommended replacement BUSCO pipeline (Simão et al. 2015). We observed more modest improvements (~10%) in the number of complete genes in the zebra finch (and no change with the hummingbird) when assessed using the BUSCO v2.0 pipeline on a set of 303 single-copy conserved eukaryotic genes (**Fig. 1B**), and barely any change (1-3%) when using a newly generated BUSCO set of 4915 avian genes (**Fig. 1C**). However, we believe that the moderate increase or no change is due to the fact that much of the BUSCO gene sets were generated from incomplete genome assemblies with short- to intermediate-length reads, particularly the 4915 protein coding avian gene set that several of the authors of the current study helped generate from the 40+ avian species in the Avian Phylogenomics Project (Zhang et al. 2014c), including the reference hummingbird (Zhang et al. 2014a). Supporting this view, we extracted the overlapping orthologous genes in the different CEGMA and BUSCO datasets, and found that the CEGMA genes are on average significantly longer than their BUSCO gene counterparts (**Suppl. Fig. 3**). When we manually examined genes randomly, we found that many of the BUSCO protein coding sequences were truncated relative to the corresponding CEGMA gene and the PacBio-based assemblies (e.g. the ribosomal protein RLP24: BUSCO (aves) is 117 a.a.; CEGMA & PacBio assembly is 163 a.a.). When compared to the CEGMA 303 eukaryotic set that includes several higher-quality genome assemblies, the PacBio-based assemblies had very few fragmented genes compared to the Sanger-based and Illumina-based assemblies (**Fig. 1B**). Thus, our new assemblies have the potential to upgrade the BUSCO set to more complete and more accurately assembled genes, a conclusion supported by our analyses below.

**Figure 1.**
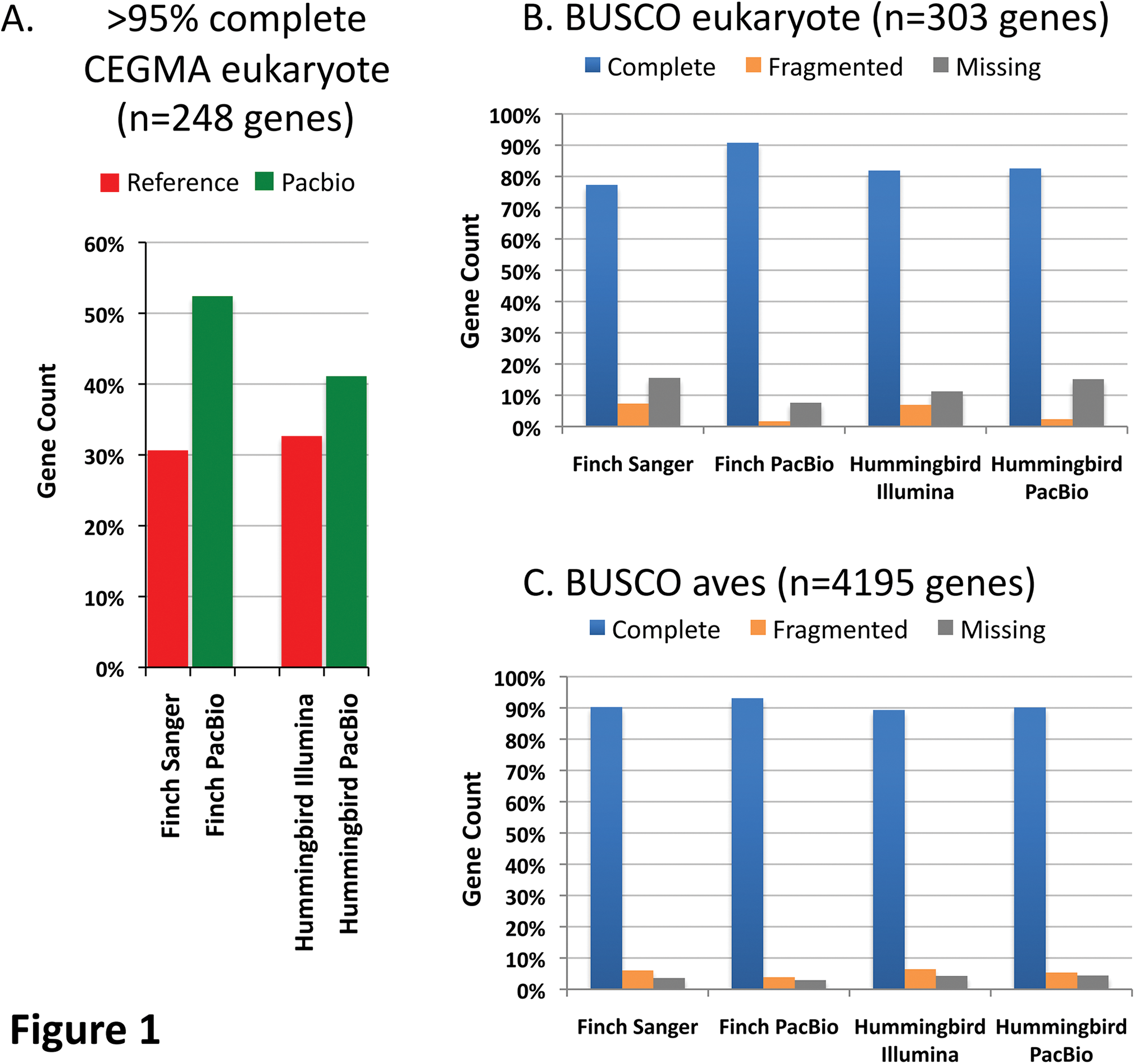
Gene completeness within assemblies. *(A)* Comparison to a 248 highly conserved core CEGMA eukaryote gene set using human genes (Parra et al. 2009), between the Sanger-based zebra finch and Illumina-based Anna’s hummingbird references and their respective PacBio-based assemblies. We used a more stringent cut-off (> 95%) for completeness than usually done (> 90%). Gent count is the percentage of genes in each of the assemblies that met this criterion. *(B)* Comparison to a 303 single-copy conserved eukaryotic BUSCO gene set (Simão et al. 2015). Complete is ≥ 95% complete; fragmented is < 95% complete; missing is not found. *(C)* Comparison to 4915 single-copy conserved genes from the avian BUSCO gene (Simão et al. 2015).

### The long-read assemblies have greater and more accurate transcriptome and regulome representations

To assess transcriptome gene completeness by an approach that does not depend on other species’ genomes, we aligned zebra finch brain paired-end Illumina RNA-Seq reads to the zebra finch genome assemblies using TopHat2 (Kim et al. 2013). We generated RNA-Seq data from microdissected RA song nucleus, a region that has convergent gene regulation with the human laryngeal motor cortex (LMC) involved in speech production (**Suppl. Fig. 4**; (Pfenning et al. 2014)). The PacBio-based assembly resulted in a ~7% increase in total transcript read mappings compared to the Sanger-based reference (**Fig. 2A**), suggesting more genic regions available for read alignments. This was explained by a decrease in unmapped reads and increase in transcripts that mapped to the genome more than once (multiple) compared to the Sanger-based reference (**Fig. 2B**), supporting the idea that the long-read assemblies recovered more repetitive or closely related gene orthologs. The PacBio assembly also resulted in ~6% more concordant aligned paired-end reads (**Fig. 2A**), indicating a more structurally accurate assembly compared to the Sanger-based reference. RNA-Seq data from the other principle brain song nuclei (HVC, LMAN, and Area X) and adjacent brain regions containing multiple cell types (**Suppl. Fig. 4A**; (Jarvis et al. 2013)) gave very similar results, with 7-11% increased mappings to the PacBio assembled genome (not shown).

**Figure 2.**
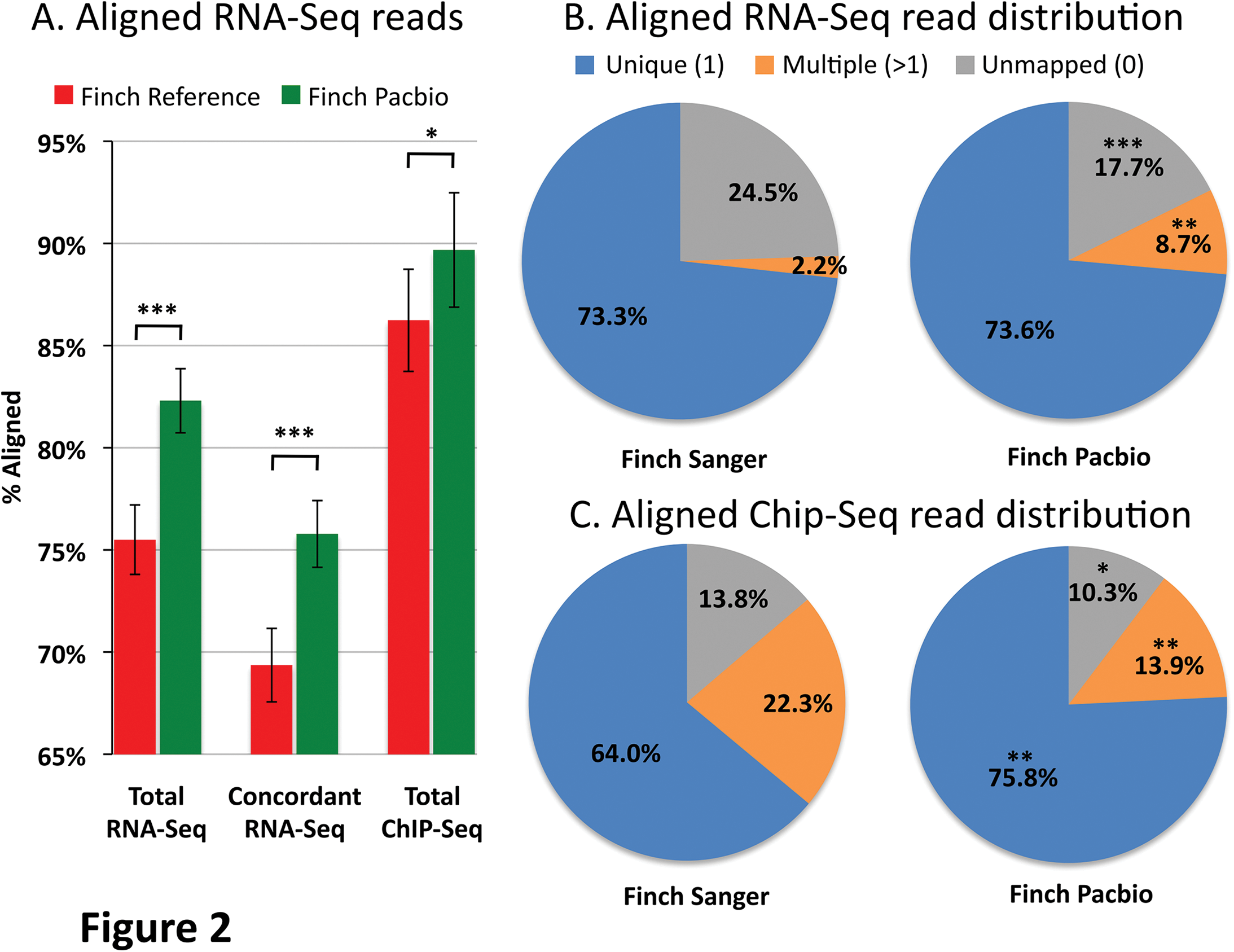
Transcriptome and regulome representation within assemblies. *(A)* Percentage of RNA-Seq and H3K27Ac ChIP-Seq reads from the zebra finch RA song nucleus mapped back to the zebra finch Sanger-based and PacBio-based genome assemblies. *(B)* Pie charts of the distributions of the RNA-Seq reads mapped to the zebra finch genome assemblies. *(C)* Pie charts of the distribution of ChIP-Seq reads mapped to the zebra finch genome assemblies. * p < 0.05; ** p < 0.002; *** p < 0.0001; paired t-test within animals between assemblies; n = 5 RNA-Seq and n = 3 ChIP-Seq independent replicates from different animals.

Regulatory regions have been difficult to identify in the zebra finch genome, as they often are GC-rich and hard to sequence and assemble with short-read technologies. To assess the regulome, we aligned HK327ac ChIP-Seq reads generated from the RA song nucleus (see methods and (Whitney et al. 2014)) to the zebra finch genome assemblies using Bowtie2 (Langmead and Salzberg 2012). H3K27ac activity is generally high in active gene regulatory regions, such as promoters and enhancers (Shlyueva et al. 2014). Similar to the transcriptome, there was an increase (~4%) of HK327ac Chip-Seq genomic reads that mapped to the PacBio-based assembly compared to the Sanger-based reference **(Fig. 2A**). Unlike the RNA-Seq transcript reads, the ChIP-Seq genomic reads showed a significant 10% increase in unique mapped reads with a concomitant decrease in multiple mapped reads (**Fig. 2B**). We believe this difference is due to technical reasons in using paired-end transcript (RNA-Seq) versus single-end genomic (ChIP-Seq) read data, as a multiple-mapped increase with the RNA-Seq transcript data was not detected when using only one read of each pair-end (*p*=0.3, paired t-test, n=5). Overall, these findings are consistent with the PacBio-based assembly having a more complete and structurally accurate assembly for both coding and regulatory non-coding genomic regions.

### Completion and correction of genes important in vocal learning and neuroscience research

The genome-wide analyses above demonstrates improvements to overall genome assembly quality using long reads, but they do not inform about real-life experiences with individual genes where there have been challenges with assemblies. We undertook a more detailed analysis of four of our favorite genes that have been highly studied in neuroscience and in particular in vocal learning/language research: *EGR1*, *DUSP1*, *FOXP2*, and *SLIT1*.

#### EGR1

The early growth response gene 1 (*EGR1*) is an immediate early gene transcription factor whose expression is regulated by activity in neurons, and is involved in learning and memory (Veyrac et al. 2014). It is up-regulated in song-learning nuclei when vocal learning birds produce learned song (Jarvis and Nottebohm 1997), along with 10% of the transcribed genome in different cell types of the brain (Whitney et al. 2014). Studying the mechanisms of regulation of *EGR1* and other immediate early genes has been an intensive area of investigation (Flavell and Greenberg 2008; Cortés-Mendoza et al. 2013), but in all intermediate- and short-read bird genome assemblies we examined thus far, part of the GC-rich promoter region is missing (**Fig. 3A, gap 1**).

**Figure 3.**
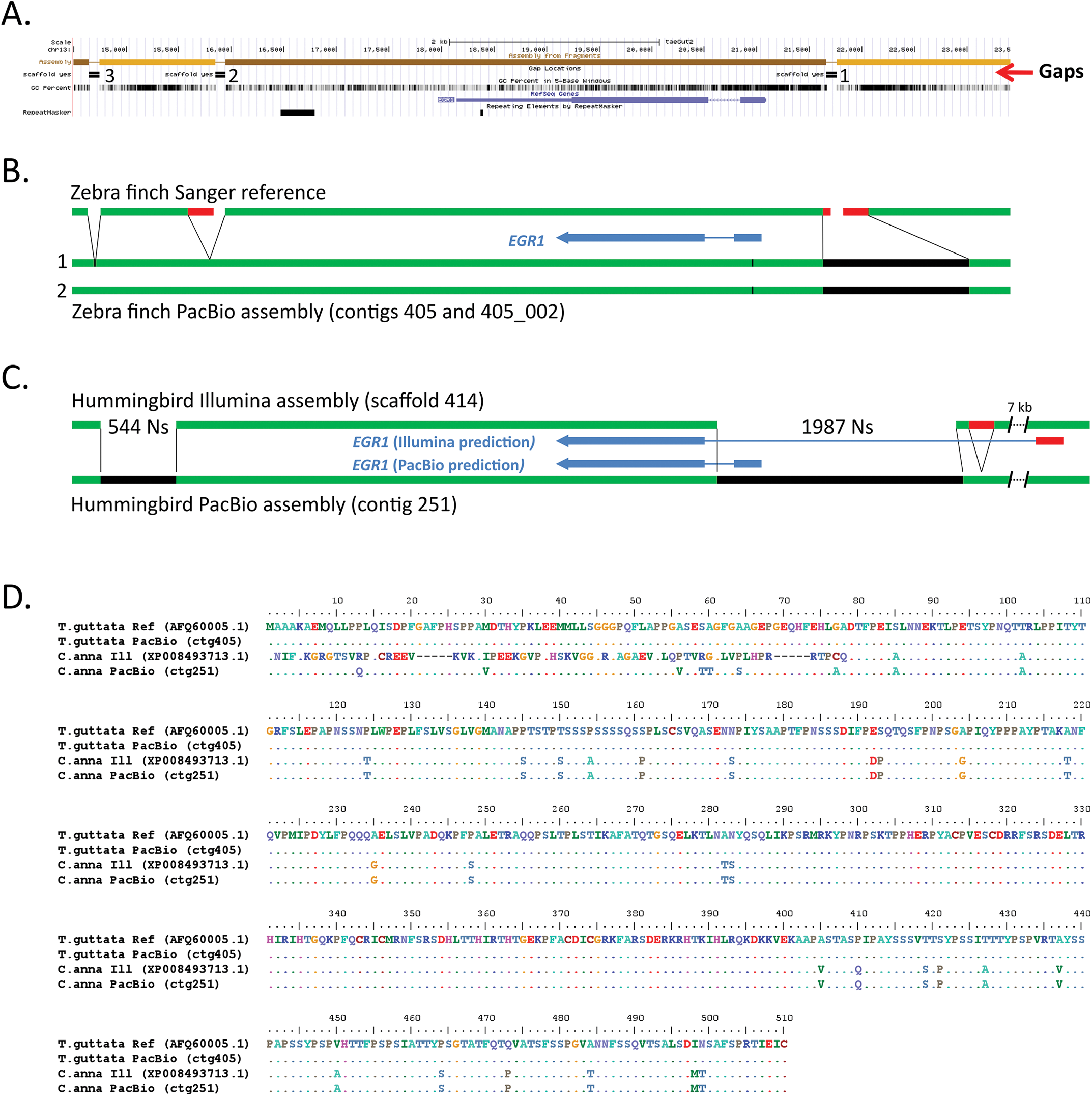
Comparison of *EGR1* assemblies. *(A)* UCSC Genome browser view of the Sanger-based zebra finch *EGR1* assembly, highlighting (from top to bottom) four contigs (light and dark brown) with three gaps, GC content, RefSeq gene prediction, and areas of repeat sequences. *(B)* Summary comparison of the Sanger-based and PacBio-based zebra finch assemblies, showing in the latter filling the gaps (black) and correcting erroneous reference sequences surrounding the gaps (red). Tick mark is a synonymous heterozygous SNP in the coding region between the primary (1) and secondary (2) haplotypes. *(C)* Comparison of the hummingbird Illumina- and PacBio-based assemblies, showing similar corrections that further lead to a correction in the protein coding sequence prediction (blue). *(D)* Multiple sequence alignment of the EGR1 protein for the four assemblies (two zebra finch and two hummingbird) in *B* and *C*, showing corrections to the Illumina-based hummingbird protein prediction by the PacBio-based assembly.

In the zebra finch Sanger-based reference, *EGR1* is located on a 5.7 kb contig (on chromosome 13), bounded by the gap in the GC-rich promoter region and 2 others downstream of the gene; gaps between contigs in the published reference were given arbitrary 100 Ns (Warren et al. 2010). We found that the PacBio long-read assembly completely resolved all three gaps in the zebra finch *EGR1* locus for both alleles, resulting in complete protein coding and surrounding gene bodies on a 205.5 kb primary contig and a 129.1 kb secondary haplotig (**Fig. 3B**; **Suppl. Fig. 5A**). The promoter region gap, located 572 bp upstream of the start of the first exon, was resolved by an 804 bp 70.1% GC-rich PacBio-based sequence (**Fig. 3B, black**). In addition to the 100 Ns in the reference, there were 241 bp to the left and right of this gap of low quality sequence (<QV40; **Fig 3B, red**) that was not supported by the PacBio data. For the second gap located ~2.2 kb downstream of the *EGR1* gene, there was an adjacent 210 bp low-similarity tandem repeat region that was not supported by the PacBio data (**Fig 3B, gap 2, red**). The third 100 N gap, located ~3.5 kb downstream of the *EGR1* gene, was resolved by 18 bp of sequence in the PacBio assembly (**Fig. 3B, gap 3**). The PacBio-based differences in the assembly were supported by numerous long-read (>10,000 bp) molecules that extended through the entire gene, spanning all three gaps (**Suppl. Fig. 6A**). The two haplotypes were >99.8% identical over the region shown (**Fig. 3B**), with only one synonymous heterozygous SNP in the coding sequence (G at position 169,283 in the primary contig 405; T at position 92,478 in secondary contig 405_002; tick mark in **Fig. 3B**).

In the Illumina-based hummingbird reference, *EGR1* was represented by 3 contigs separated by 2 large gaps of 544 Ns and 1987 Ns respectively (**Fig. 3C**), in a large 2.98 Mb scaffold. In contrast, in the PacBio-based hummingbird assembly, *EGR1* was fully resolved in a large 810 kb contig (**Fig. 3C**). Gene prediction (using Augustus (Stanke et al. 2008)) yielded a protein of the same length as the finch EGR1 protein (510 a.a.), and with high (93%) sequence homology (**Fig. 3D**). The PacBio-based assembly revealed that the larger gap in the Illumina-based assembly harbors the beginning of the *EGR1* gene, including the entire first exon, two thirds of the first intron, and the GC-rich promoter region (**Fig. 3C, black**). Due to this gap in the reference, the corresponding NCBI gene prediction (accession XP_008493713.1) instead recruited a stretch of sequence ~7 kb upstream of the gap, predicting a first exon that has no sequence homology with *EGR1* in the PacBio-based assembly or to sequences of other species (**Fig. 3C & D**). Upstream of this gap in the Illumina-based assembly was also a 200 bp tandem repeat that was not supported by the PacBio sequence reads and the assembly (**Fig. 3C, red; Suppl. Fig. 5B**). These PacBio-based differences in the assembly were further validated by single-molecule Iso-Seq mRNA long-reads of *EGR1* from a closely related species (the Ruby-throated hummingbird; kindly provided by R. Workman & W. Timp) that fully contained both predicted exons (**Suppl. Fig. 6B**). The PacBio-based assembly did not generate a secondary haplotype for this region, indicating that the two alleles are identical or nearly identical for the entire 810 kb contig in the individual sequenced. Upstream and downstream of a high homology region that includes the *EGR1* exons, intron, and GC-rich promoter, there was little sequence homology between the PacBio-based hummingbird and zebra finch assemblies (**Suppl. Fig. 7**).

These findings indicate that relative to the intermediate- and short-read assemblies, the PacBio-based long-read assembly can fill in missing gaps in a previously hard-to-sequence GC-rich regulatory region, eliminate low quality erroneous sequences and base calls at the edges of gaps in a Sanger-based assembly, and eliminate erroneous tandem duplications adjacent to gaps, all preventing inaccurate gene predictions. In addition, using one species as a reference to help assemble another may not work for such a gene, as the surrounding sequence to the gene body in these two Neoaves species is highly divergent.

#### DUSP1

The dual specificity phosphatase 1 (*DUSP1*) is also an immediate early gene, but one that regulates the cellular responses to stress (Liu et al. 2008). In all species examined thus far it is mostly up-regulated by activity in the highly active thalamic-recipient primary sensory neurons of the cortex (i.e. mammal cortex layer 4 cells and the comparable avian intercalated pallial cells), but within the motor pathways, it is only up-regulated to high levels by activity in the vocal learning circuits of vocal learners (Horita et al. 2010; Horita et al. 2012). This specialized regulation in vocal learning circuits has been proposed to be associated with convergent microsatellite sequences found in the upstream promoter region of the gene mainly in vocal learning species (Horita et al. 2012). This was determined by PCR-cloning of single genomic molecules of multiple species, because the reference assemblies did not have this region properly assembled (Horita et al. 2012).

In the zebra finch Sanger-based reference, *DUSP1* is located on the chromosome 13 scaffold, separated in 3 contigs, with 2 gaps (**Fig. 4A**). The NCBI gene prediction of this assembly resulted in 4 exons generating a 322 a.a. (XP_002192168.1), which is ~13% shorter than the *DUSP1* homologs of other species, e.g. chicken (369 a.a., Genbank accession NP_001078828), rat (367 a.a., NP_446221), and human (367 a.a, NP_004408). The 2 gaps coincide with the end of the first predicted exon and the beginning of the third predicted exon (**Fig. 4A**). An additional gap upstream of the coding sequence falls within the known microsatellite repeat region (**Fig. 4A**). The PacBio-based assembly completely resolved the entire region for both alleles, in an 8.4 Mb primary contig and an 8.0 Mb secondary haplotig (**Fig. 4B, Suppl. Fig. 8A**). The Augustus gene prediction resulted in a protein with 4 exons but now with a total length of 369 a.a. that was homologous across its length to *DUSP1* of other vertebrate species (e.g., 96% with chicken GGv5 assembly, also recently updated with long reads). Comparing the two assemblies revealed that: 1) the first exon in the Sanger-based reference is truncated by 28 a.a. in the gap; 2) within a 20 a.a. region near the edge of that truncation are three a.a. different from the reference (**Fig. 4**; residues 81, 89, and 98), but the same as other songbird species (**Suppl. Fig. 9C**) and with strong support in the PacBio reads (**Suppl. Fig. 9D**); 3) the second exon and adjacent intron is missing a 80.8% GC-rich 0.46 kb sequence in the reference, and is instead replaced by a 1.7 kb contig of a partially repeated sequence from the microsatellite region upstream of *DUSP1* (R’ in **Fig. 4B**), part of which was erroneously recruited in the second exon of the NCBI reference gene prediction (**Fig. 4D**); and 4) the microsatellite repeat itself is erroneously partially duplicated in the reference, flanking both sides of gap 1 (R’’ in **Fig. 4B**). Our PacBio phased assembly revealed why both instances of R’ are not identical in the reference, because they in fact belong to the different haplotypes: the 1.7 kb contig corresponds to the upstream region in the primary PacBio haplotype (contig 32) whereas the actual upstream region in the reference corresponds to the upstream region in the secondary PacBio haplotype (contig 32_022) (**Fig. 4B**). This main microsatellite region is 76 bp longer (796 *vs.* 720 bp) in the primary haplotype, and the neighboring smaller upstream microsatellite contains 3 additional 20-21 bp repeats (11 *vs.* 8) in the primary haplotype (**Suppl. Fig. 9A**). Within the protein coding sequence there were four synonymous heterozygous SNPs between haplotypes (not shown).

**Figure 4.**
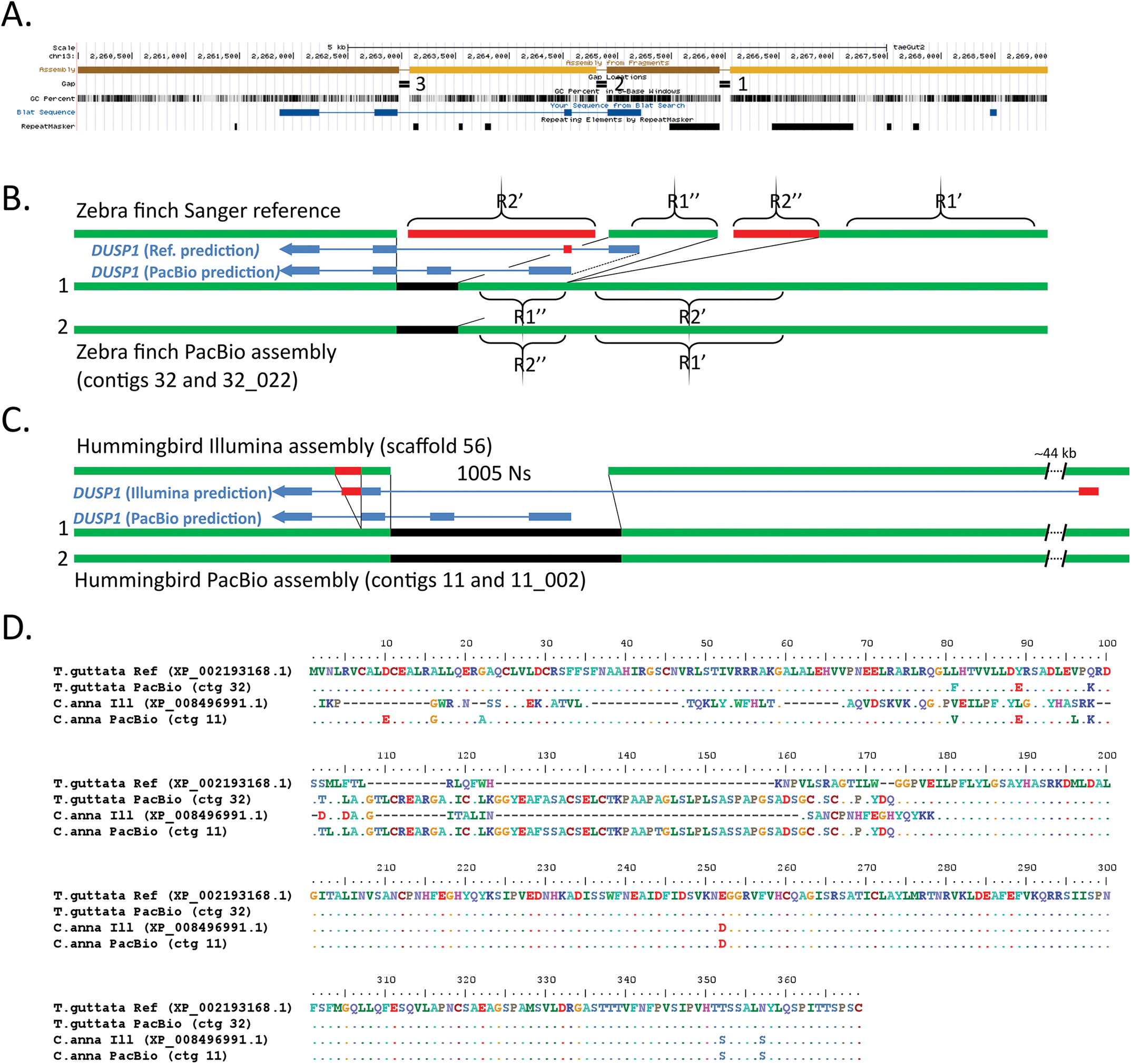
Comparison of *DUSP1* assemblies. *(A)* UCSC Genome browser view of the Sanger-based zebra finch *DUSP1* assembly, highlighting four contigs with three gaps, GC content, Blat alignment of the NCBI gene prediction (XP_002193168.1, blue), and repeat sequences. *(B)* Resolution of the region by the PacBio-based zebra finch assembly, filling the gaps (black) and correcting erroneous reference sequences in repeat regions (red) and gene predictions (blue). *(C)* Resolution and correction to the hummingbird Illumina-based assembly with the PacBio-based assembly (same color scheme as in *B*). *(D)* Multiple sequence alignment of the DUSP1 protein for the four assemblies in *B* and *C*, showing numerous corrections to the Sanger-based and Ilumina-based protein predictions by both PacBio-based assemblies.

In the hummingbird Illumina-based assembly, the *DUSP1* region was represented by 2 contigs separated by a large 1005 N gap (**Fig. 4C**), on a 7 Mb scaffold. In the PacBio-based assembly, the entire gene was fully resolved (**Fig. 4C; Suppl. Fig. 8B**), in a much larger gapless 12.8 Mb contig (the second allele is fully resolved in a 3.8 Mb contig). Comparing the two assemblies revealed that because of the gap in the Illumina-based reference, it lacks about half of the *DUSP1* gene, including the first two exons and introns, and ~380 bp upstream of the start of the gene (**Fig. 4C**). As a result, the corresponding NCBI gene prediction (XP_008496991.1) recruited a sequence ~44 kb upstream predicting 46 a.a. with no sequence homology to *DUSP1* of other species, whereas the PacBio-based assembly yielded a 369 a.a. protein with 99% sequence homology to the PacBio-based zebra finch and chicken *DUSP1* (**Fig. 4D**). A 200 bp tandem repeat in the Illumina-based assembly downstream of the gap, erroneously in exon 3, is a misplaced copy of the microsatellite region (**Fig. 4C; Suppl. Fig. 8B**). This is the reason why two thirds of exon 3 is erroneously duplicated in the NCBI protein prediction (**Fig. 4D**). The PacBio assemblies also revealed that the microsatellite region was significantly shorter in the hummingbird (~270 bp) than the zebra finch genome (~1100 bp; **Suppl. Fig. 9B**).

These findings in both species demonstrate that intermediate- and short-read assemblies not only have gaps with missing relevant repetitive microsatellite sequence, but that short-read misassemblies of these repetitive sequences lead to erroneous protein coding sequence predictions. Further, not only does the long-read assembly resolve them, but it helps generate a diploid assembly that reveals allelic differences, and prevents erroneous assembly duplications and misplacement errors.

#### FOXP2

The forkhead box P2 (*FOXP2)* gene plays an important role in spoken-language acquisition (Fisher and Scharff 2009). In humans, a point mutation in the protein coding binding domain in the KE family (Lai et al. 2001) as well as deletions in the non-coding region of *FOXP2* (Turner et al. 2013) results in severe spoken language impairments in heterozygous individuals (homozygous is lethal). In songbirds, FOXP2 expression in the Area X song nucleus is differentially regulated by singing activity and during the song learning critical period, and is necessary to properly imitate song (Haesler et al. 2004; Teramitsu and White 2006; Haesler et al. 2007). In mice, although vocalizations are mainly innate, animals with the KE mutation demonstrate a syntax apraxia-like deficit in syllable sequencing similar to that of humans (Castellucci et al. 2016; Chabout et al. 2016). Thus, *FOXP2* has become the most studied gene for understanding the genetic mechanisms and evolution of spoken language (Condro and White 2014), yet we find that the very large gene body of ~400 kb is incompletely assembled, including in vocal learning species (**Fig. 5A**).

**Figure 5.**
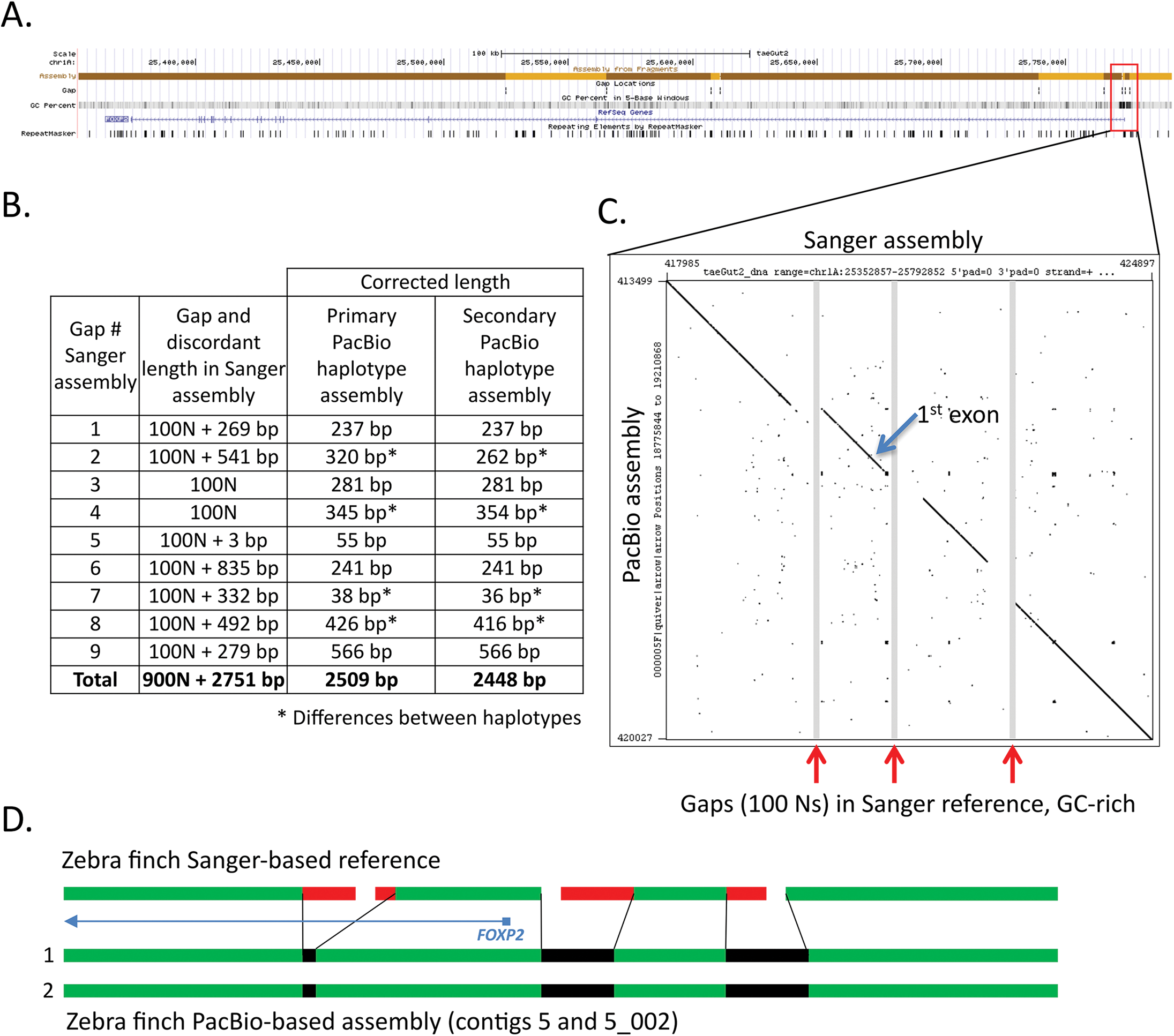
Comparison of *FOXP2* assemblies. *(A)* UCSC Genome browser view of the Sanger-based zebra finch *FOXP2* assembly, highlighting 10 contigs with 9 gaps, GC content, RefSeq gene prediction, and repeat sequences. *(B)* Table showing the number of resolved and corrected erroneous base pairs in the gaps by the PacBio-based primary and secondary haplotype assemblies; * indicates differences between haplotypes. *(C)* Dot plot of the Sanger-based reference (x-axis) and the PacBio-based primary assembly (y-axis) corresponding to the three GC-rich region gaps immediately upstream and surrounding the first exon of the *FOXP2* gene. *(D)* Schematic summary of corrections to the three gaps shown in *C*, in the two haplotypes of the PacBio-based assembly. The protein coding sequence alignments are in Supplementary Figure 11A.

In the zebra finch Sanger-based reference, *FOXP2* is located on the chromosome 1A scaffold and separated into 10 contigs (1 to 231 kb in length) with nine 100 N gaps each (**Fig. 5A**). These include 2 gaps immediately upstream of the first exon, making the beginning of the gene poorly resolved. The provisional RefSeq mRNA for *FOXP2* (NM_001048263.1) contains 19 exons and encodes a 711 a.a. protein (NP_001041728.1). In the PacBio-based assembly, the entire 400 kb gene is fully resolved for both haplotypes in 21.5 Mb and 7.6 Mb contigs, respectively (**Suppl. Fig. 10A**). As observed in the previous examples, sequences of various sizes surrounding all 9 gaps in the Sanger-based reference were unsupported by the PacBio data, resulting in a total of 2509 bp of corrected sequence in the PacBio-based primary haplotype (**Fig. 5B**). The two filled gaps in the upstream region and the next gap in the first intron were GC-rich (77.6%, 66.5%, and 67.8%, respectively) (**Fig. 5A,C**), indicative of the likely cause of the poor quality Sanger-based reads (**Fig. 5D**). The DNA sequence between the two assembled PacBio haplotypes was >99% similar across the entire 400 kb *FOXP2* gene, and identical over the coding sequence, with differences occurring in the more complex non-coding gaps that were difficult to sequence and assemble by the Sanger method (**Fig. 5B \***61 nucleotide differences total). The predicted protein sequence from the PacBio-based assembly is identical to the predicted Sanger-based reference (NP_001041728.1), with the exception of a.a. residue 42 (threonine *vs.* serine) (**Suppl. Fig. 11A**). The PacBio nucleotide call also exists in the mRNA sequence of another zebra finch animal in NCBI (NM_001048263.2) and in other avian species we examined, and is thus likely a base call error in the Sanger-based zebra finch reference.

In the hummingbird Illumina-based assembly, as expected with short-read assemblies relative to the Sanger-based zebra finch reference, the *FOXP2* gene was even more fragmented, in 23 contigs (ranging 0.025 to 2.28 kb in lengths) with 22 gaps (**Suppl. Fig. 10B**). The two largest gaps encompass the beginning of the gene and first (non-coding) exon, resulting in corresponding low quality predicted mRNA (XM_008496149.1). The predicted protein (XP_008494371.1) includes an introduced correction (a.a. 402; **Suppl. Fig. 11A**, X nucleotide) to account for a genomic stop codon, and an 88 N gap within exon 6 that artificially splits the exon into two pieces (**Suppl. Fig. 11B**). In the hummingbird PacBio-based assembly, the *FOXP2* gene is fully resolved and phased into two haplotype contigs of 3.2 Mb each (**Suppl. Fig. 10B**). The erroneous stop codon is corrected (2170128C (ctg 110) and 2183088C (ctg 110_009), instead of 841788T (Illumina assembly scaffold 125)), and exon 6 is accurately contiguous, removing the gap and an additional 22 bp of erroneous tandem repeat sequence adjacent to the gap (**Suppl. Fig. 11B & C**). The PacBio-based assembly also corrects three other instances of erroneous tandem duplications over the gene region in the Illumina-based assembly, as well as removes a 462 bp stretch of sequence adjacent to a long homonucleotide A stretch in intron 1 of the Illumina-based assembly (position 972040) (**Suppl. Fig. 12A**). The two PacBio assembled haplotypes are >99% similar, with one heterozygous SNP (2172601T (contig 110) *vs.* 2185560A (contig 110_009)) in exon 6 that is silent, and a 708 bp deletion in the secondary haplotype (contig 110_009 (at position 2128952) relative to contig 110; **Suppl. Fig. 12B**). The Illumina-based assembly has the deleted allele.

These findings replicate those of the previously discussed genes, and in addition show that the PacBio-based assembly can fully resolve very large genes and erroneous assembled sequence in gaps due to repeats or homonucleotide stretches, and reveal large haplotype differences. The phased, diploid assembly avoids the possibility of large missed sequences due to deletion in one allele.

#### SLIT1

Slit homolog 1 (*SLIT1*) is a repulsive axon guidance ligand for the *ROBO1* receptor, and is involved in circuit formation in the developing brain (Blockus and Chédotal 2014). Recently, *SLIT1* was shown to have convergent specialized down-regulated expression compared to the surrounding brain region in the RA song nucleus of all independently evolved vocal learning bird lineages and in the analogous human LMC (Pfenning et al. 2014; Wang et al. 2014) (**Suppl. Fig. 4**), indicating a potential role of *SLIT1* in the evolution and formation of vocal learning brain circuits. A fully resolved *SLIT1*, including regulatory regions, is necessary to assess the mechanisms of its specialized regulation in vocal learning brain regions.

In the zebra finch Sanger-based reference, *SLIT1* is located on chromosome 6, split among 8 contigs with 7 gaps, and 7 additional contigs and gaps surrounding the ~40 kb gene (**Fig. 6A**). The SLIT1 gene is complex, with over 35 exons. We noted an incomplete predicted protein of the reference (XP_012430014.1) relative to some other species (chicken (NM_001277336.1), human (NM_003061.2), and mouse (NM_015748.3)), and our *de novo* gene predictions from the reference also resulted in a truncated protein with two missing exons (**Fig. 6B**). The PacBio-based assembly fully resolved the gene region, in two alleles on 15.7 Mb and 5.6 Mb contigs, respectively, and completely recovered all 35+ exons (**Suppl. Fig. 13A**). Similar to above, reference sequences flanking the gaps were found to be erroneous and corrected, and an erroneous tandem duplication was also corrected (not shown). Filling in these gaps recovered the two missing exons: exon 1 within a 1 kb region of sequence in the PacBio-based assembly that is 75% GC-rich, replacing 390 bp of erroneous gap-flanking sequence, and exon 35 adjacent to a gap (**Fig. 6A & B**). The PacBio-based assembly thereby generates a complete *SLIT1* gene prediction of 1538 a.a. (**Fig. 6B**). The gene is heterozygous in the individual, with 3 codon differences between the two alleles (**Fig. 6B**, positions 90, 1006, and 1363, respectively), and an additional 24 silent heterozygous SNPs across the coding region. The two alleles were phased along the entire length of the gene.

**Figure 6.**
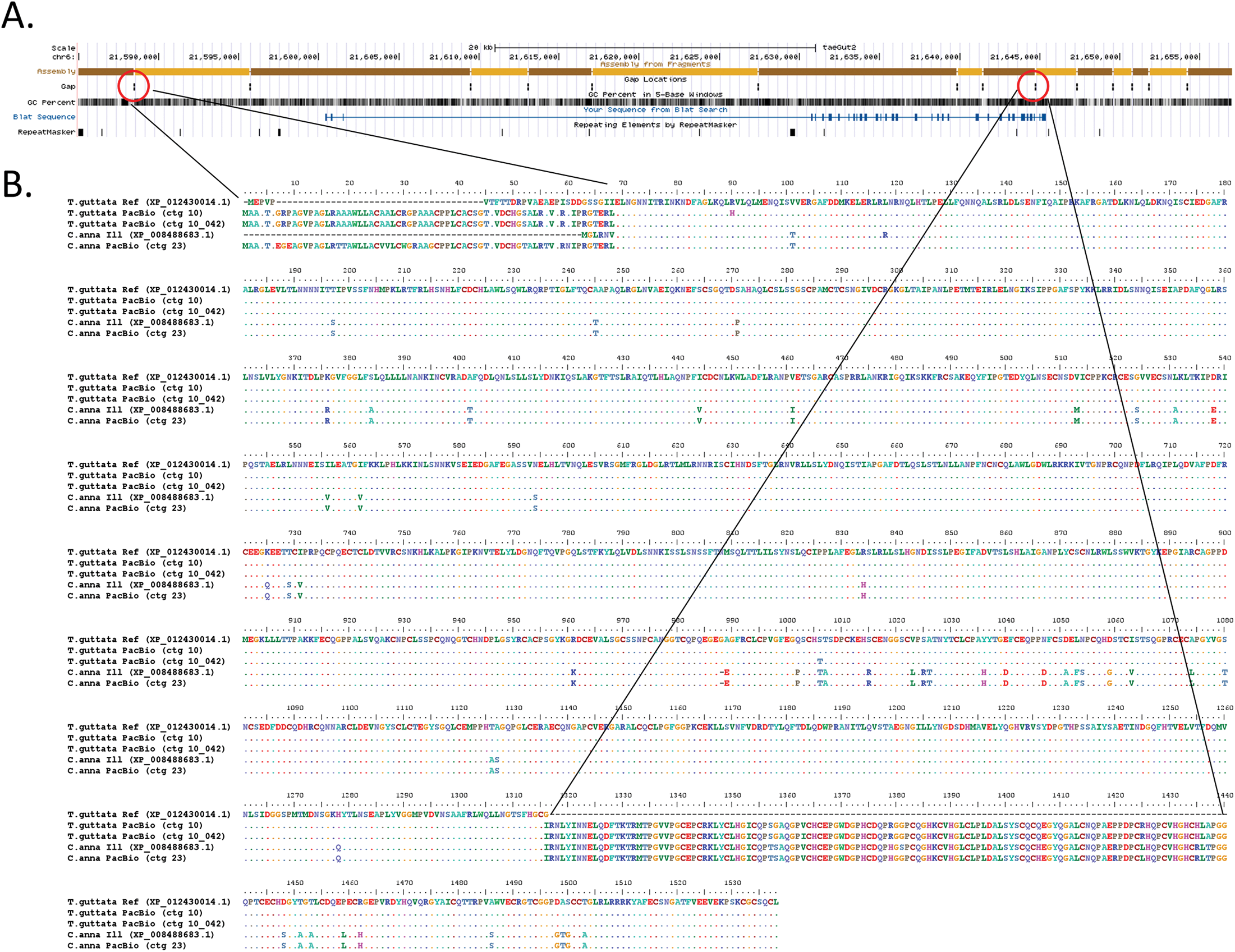
Comparison of *SLIT1* assemblies. *(A)* UCSC Genome browser view of the Sanger-based zebra finch *SLIT1* assembly, highlighting 15 contigs with 14 gaps, GC content, NCBI *SLIT1* gene prediction (XP_012430014.1, blue), and repeat sequences. Red circles, gaps that correspond to the missing exon 1 and part of the missing exon 35, respectively. *(B)* Multiple sequence alignment comparison of the SLIT1 protein for four assemblies compared, including the two different haplotypes from the PacBio-based zebra finch assembly (rows 2 and 3).

In the hummingbird Illumina-based assembly, the *SLIT1* gene is separated on 9 contigs with 8 gaps ranging in length from 91 to 1018 bp, comprising 3320 bp of missing sequence, or 5.3% of the gene region (**Suppl. Fig. 13B**). The PacBio-based assembly fully resolved and phased *SLIT1* into haplotypes on 9.9 Mb contigs (**Suppl. Fig. 13B**). The resulting protein of 1538 a.a. has high homology to the zebra finch PacBio-based *SLIT1* (95% a.a. identity; **Fig. 6B**) and the individual is homozygous for the SLIT1 protein. Comparisons revealed that as with the Sanger-based reference, the first exon (68 a.a.) is missing completely in the Illumina-based assembly (**Fig. 6B**), corresponding to a gap of 495 Ns, which the PacBio-based assembly replaced by a 567 bp 76% GC-rich sequence (**Suppl. Fig. 13B**). In addition, there were two sequence errors in the Illumina-based assembly, which resulted in erroneous amino acid predictions in the SLIT1 protein (**Fig. 6B**, positions 118 and 1381, respectively).

These findings demonstrate that long-read assemblies can fully resolve a complex multi-exon gene, as well as have a higher base-call accuracy than Sanger- or Illumina-based reads in difficult to sequence regions, including exons, leading to higher protein-coding sequence accuracy.

#### Other genes

We have manually compared several dozen other genes between the different assemblies, and found in all cases investigated that errors in the Sanger-based and Illumina-based assemblies were corrected in the PacBio-based long-read assemblies. These genes included other immediate early gene transcription factors, other genes in the *SLIT* and *ROBO* gene families, and the *SAP30* gene family, which all had the same types of errors in the genes discussed above. In addition, we also found cases were genes were missing from the Sanger-based zebra finch or Illumina-based hummingbird assemblies entirely, and could have been interpreted as missing in these species. These included the DNA methyltransferase enzyme *DNMT3A* missing in the Sanger-based finch assembly and *DRD4* missing in the hummingbird assembly (Haug-Baltzell et al. 2015), with both fully represented in the PacBio-based assemblies. We also noted cases where an assembled gene was incorrectly localized on a scaffold in the Sanger-based assembly whose synteny was corrected with the PacBio-based assembly, such as the vasopressin receptor AVPR1B, which will be reported on in more detail separately. Data for these types of errors were not shown due to space limitations, but they offer further examples of the important improvements of PacBio long-read technology for generating more accurate genome assemblies.

## Conclusions

Although the intermediate-read and short-read assemblies had correct sequences and assembled regions in terms of total base pairs covered, the long-read assemblies revealed numerous errors within and surrounding many genes. These errors are not simply in so-called “junk” intergenic repetitive DNA known to be hard to assemble with short reads (Treangen and Salzberg 2011; Palazzo and Gregory 2014), but within functional regions of genes. The assemblers for the short reads sometimes take a repetitive sequence, some in functional repetitive regulatory regions, and insert them in a non-repetitive region of a gene, resulting in an error. Some of these assembly errors and gaps in the sequences lead to gene and protein coding sequence prediction errors, sometimes recruiting completely wrong sequence in the protein. The PacBio-based long-read assemblies corrected these problems, and for the first time resolved gene bodies of all the genes we examined into single, contiguous, gap-less sequences. The phasing of haplotypes, although initially done to prevent a computationally introduced indel error, reveal how important phasing is to prevent assembly and gene prediction errors. Thus far, we have not seen an error (i.e. difference) in the genes we examined in the PacBio-based long-read assembly relative to the other assemblies that was supported by single sequenced genomic DNA molecules, RNA-Seq and Iso-Seq mRNA molecules, or other independent evidence. With these improvements, we now, for the first time, have complete and accurate assembled genes of interest that we now can pursue further without the need to individually and arduously clone, sequence, and correct the assemblies one gene at a time.

Our study highlights the value of maintaining frozen tissue or cells of the individuals used to create previous reference genomes, as we could only discover some of the errors (e.g. caused by haplotype differences) by long-read *de novo* genome assemblies of the same individual used to create the reference. We are now using these PacBio-based assemblies with several groups and companies as starting assemblies for scaffolding into phased, diploid, chromosome-level zebra finch and hummingbird assemblies to upgrade the references, which will be reported on separately. However, even without scaffolding, these more highly contiguous assemblies will be helpful to researchers to extract more accurate assemblies of their genes of interests, saving a great amount of time and energy, while adding new knowledge and biological insights.

## Materials & Methods

### DNA isolation

For both the zebra finch and hummingbird, frozen muscle tissue from the same animals used to create the Sanger-based (Warren et al. 2010) and Illumina-based (Zhang et al. 2014c) references, respectively, was processed for DNA isolation using the KingFisher Cell and Tissue DNA Kit (97030196). Tissue was homogenized in 1 ml of lysis buffer in M tubes (Miltenyi Biotec) using the gentleMACS™ Dissociator at the Brain 2.01 setting for 1 minute. The cell lysate was treated with 40 ul of protease K (20mg/ml) and incubated overnight. DNA was purified using the KingFisher Duo system (5400100) using the built in KFDuoC_T24 DW program.

### Library preparation and sequencing

For the zebra finch, two samples were used for library construction. Each DNA sample was mechanically sheared to 60 kb using the Megaruptor system (Diagenode). Then >30 kb libraries were created using the SMRTbell Template Prep Kit 1.0 (Pacific Biosciences), which includes a DNA Damage Repair step after size selection. Size selection was made for 15 kb for the first sample and 20 kb for the second sample, using a Blue Pippin instrument (Sage Science) according to the protocol “Procedure & Checklist – 20 kb Template Preparation Using BluePippin Size-Selection System”. For the hummingbird, 70 ug of input DNA was mechanically sheared to 35 and 40 kb using the Megaruptor system, a SMRTbell library constructed, and size selected to > 17 kb with the BluePippin. Library quality and quantity were assessed using the Pippin Pulse field inversion gel electrophoresis system (Sage Science), as well as with the dsDNA Broad Range Assay kit and Qubit Fluorometer (Thermo Fisher).

SMRT sequencing was performed on the Pacific Biosciences RS II instrument at Pacific Biosciences using an on plate concentration of 125 pM, P6-C4 sequencing chemistry, with magnetic bead loading, and 360 minute movies. A total of 124 SMRT Cells were run for the zebra finch and 63 SMRT Cells for the hummingbird. Sequence coverage for the zebra finch was ~96 fold, with half of the 114 Gb of data contained in reads longer than 19 kb. For the hummingbird, coverage was ~70 fold, with half of the 40.4 Gb of data contained in reads longer than 22 kb (**Suppl. Fig. 2**).

### Assembly

Assemblies were carried out using FALCON v0.4.0 followed by the FALCON-Unzip module (Chin et al. 2016). FALCON is based on a hierarchical genome assembly process (Chin et al. 2013). It constructs a string graph from error-corrected PacBio reads that contains ‘haplotype-fused’ genomic regions as well as “bubbles” that capture divergent haplotypes from homologous genomic regions. The FALCON-Unzip module then assigns reads to haplotypes using heterozygous SNP variants identified in the FALCON assembly to generate phased contigs corresponding to the two alleles. The diploid nature of the genome is thereby captured in the assembly by a set of primary contigs with divergent haplotypes represented by a set of additional contigs called haplotigs. Genomic regions with low heterozygosity are represented as collaped haplotypes in the primary contigs. Genome assemblies were run on an SGE-managed cluster using up to 30 nodes, where each node has 512 Gb of RAM distributed over 64 slots. The same configuration files were used for both species (**Supplementary note A**). Three rounds of contig polishing were performed. For the first round, as part of the FALCON-Unzip pipeline, primary contigs and secondary haplotigs were polished using haplotype-phased reads and the Quiver consensus caller. For the second and third rounds of polishing, using the “resequencing” pipeline in SMRTlink v3.1, primary contigs and haplotigs were concatenated into a single reference and BLASR was used to map all raw reads back to the assembly, followed by consensus calling with Arrow. The raw sequence data for the hummingbird are available in Genbank (SRX1131887, SRX1130526, SRX1130525), both assemblies (BioProject ID PRJNA368994, WGS accessions pending) can be found here until they have NCBI accession numbers:

https://www.dropbox.com/s/5aw4ju5khjtpue3/Taeniopygia_guttata_PacBio_FALCON_unzip_primary_alternate_contigs.fasta.gz?dl=0

https://www.dropbox.com/s/a4sw394sghtmvsl/Calypte_anna_PacBio_FALCON_unzip_primary_alternate_haplotype.fasta.gz?dl=0

### Genome completeness

To assess quality and completeness of the assemblies, we used a set of 248 highly conserved eukaryotic genes from the CEGMA human set (Parra et al. 2009) and located them in each of the assemblies compared in this study. Briefly, the CEGMA human peptides were aligned to each genome using genblastA (She et al. 2009). The regions showing homology were then used to build gene models with exonerate (Slater and Birney 2005) which were then assessed for frameshifts using custom shell scripts. In addition, we queried each genome for a set of 303 eukaryotic conserved single-copy genes as well as from 4915 conserved single-copy genes from 40 different avian species using the BUSCOv2.0 pipeline (Simão et al. 2015).

To compare protein amino acid sequence size between the CEGMA and BUSCO datasets, we performed blastp of each CEGMA sequence against the ancestral proteins of the target BUSCO dataset. We took the single best hit with an e-value cut off of 0.001 and extracted the CEGMA and BUSCO protein lengths values. We then ran a one-sided paired Wilcoxon signed-rank test of the two lengths for each protein.

### Gene prediction

Gene predictions for the zebra finch PacBio-based assembly were conducted by running Augustus (Stanke et al. 2008) gene prediction software (v3.2.2) on the contigs, and incorporating the Illumina short read RNA-Seq brain data aligned with Tophat2 (v2.0.14, (Kim et al. 2013)) as hints for possible gene structures. The data consisted of 146,126,838 paired-end reads with an average base quality score of 36. Augustus produces a distribution of possible gene models for a given locus and models that are supported by our RNA-Seq data are given a “bonus” while the gene models not supported by RNA-Seq data are given a “penalty”. This results in the gene model most informed by biological data being selected as the most likely gene model for that locus.

We did not have Illumina transcriptome data for Anna’s hummingbird, so standard Augustus gene prediction (v3.2.2) was used with both chicken and human training background to determine the sequence predictions of the genes examined. The human-based predictions captured more of the divergent 5’ ends of the longer genes (*SLIT1* and *FOXP2*) then the chicken-based predictions, so a combination of both were used to produce the final sequences in this manuscript.

### RNA-Seq

RNA sequencing was centered around vocal learning brain regions in the zebra finch and will be described in more detail in a later publication. We utilized our data here for population analyses of assembly quality and for initial annotations. In brief, following modifications of a previously described protocol (Whitney et al. 2014), nine adult male zebra finches were isolated in soundproof chambers for 12 hours in the dark to obtain brain tissue from silent animals. Then brains were dissected from the skull and sectioned to 400 microns using a Stoelting tissue slicer (51415). The sections were moved to a petri dish containing cold PBS with proteinase inhibitor cocktail (11697498001). Under a dissecting microscope (Olympus MVX10), the four principle song nuclei (Area X, LMAN, HVC, and RA) as well as their immediate adjacent brain regions were microdissected using 2mm fine scissors and placed in microcentrifuge tubes. The samples were stored at -80 °C. Then RNA was isolated and quantified, and samples of two birds were then pooled for each replicate, resulting in 5 replicates (one single animal in one). RNA was converted to cDNA and library preparation was performed using the TruSeq Library Prep Kit V2 (Illumina) and paired-end reads were sequenced on an Illumina HiSeq 2500 system. Adapters and poor quality bases (<30) were trimmed using fastq-mcf from the ea-utilities package, and reads were aligned to assemblies using Tophat2 (v2.0.14).

### Chip-Seq

Three adult male zebra finches were treated as above, the brains dissected, and the RA and surrounding arcopallium of each bird was then processed individually using the native ChIP protocol described in (Brind’Amour et al. 2015) with an H3K27ac antibody (Ab#4729). The DNA libraries were prepared using the MicroPlex Library Preparation Kit v2 (C05010012). 50 bp single-end sequencing was done on the Illumina HiSeq 4000 system. The reads were aligned to the assemblies using Bowtie2 (v2.2.9, (Langmead and Salzberg 2012)). More detail will be provided in a later publication focusing on vocal learning brain regions.

## Acknowledgements

We thank Art Arnold for sending us the carcass with tissue used to create the original Sanger-based zebra finch assembly, and Claudio Mello and Peter Lovell again for their previous capture of the hummingbird used for the reference assembly. We thank the PacBio sequencing (Christine Lambert, Jill Muehling, Primo Baybayan) and assembly (Matthew Seetin, Richard Hall, Jane Landolin, Lawrence Hon) teams for help with sequencing and assembly, and Winston Timp and Rachael Workman for early access to their Iso-Seq reads of the Ruby-throated hummingbird. We thank Alejandro Burga for noticing indel errors in the original PacBio haploid assembly that lead us work with others to find a correction with phasing, and several members of the Jarvis lab (Mathew Biegler, Ha Na Choe, and Constantina Theofanopoulou) and Asher Haug-Baltzell for help with manual analyses of gene models between the assemblies. Finally, we thank members of the G10K and B10K consortiums for valuable discussions on metrics of genome assembly quality. This work was supported by HHMI funds to E.D.J. and PacBio funds to J. K.

## Author contributions

J.K. and E.D.J. designed the project and wrote the manuscript;C.S.C. and S.K. carried out genome assemblies; J.K., G.G. and S.K. conducted analyses on single genes as well as CEGMA and BUSCO analyses; G.G. conducted RNA-Seq analyses, L.C. conducted Chip-Seq analyses; J.H. processed samples; and all authors contributed to writing and editing the manuscript.

## Abbreviations

A1-L4: primary auditory cortex – layer 4
Am: nucleus ambiguous
Area X: a vocal nucleus in the striatum
aSt: anterior striatum vocal region
aT: anterior thalamus speech area
Av: avalanche
aDLM: anterior dorsolateral nucleus of the thalamus
DM: dorsal medial nucleus of the midbrain
HVC: a vocal nucleus (no abbreviation)
L2: auditory area similar to human cortex layer 4
LSC: laryngeal somatosensory cortex
LMC: laryngeal motor cortex
MAN: magnocellular nucleus of the anterior nidopallium
MO: oval nucleus of the anterior mesopallium
NIf: interfacial nucleus of the nidopallium
PAG: peri-aqueductal gray
RA: robust nucleus of the arcopallium
v: ventricle space

---

**Supplementary Figure 1.**
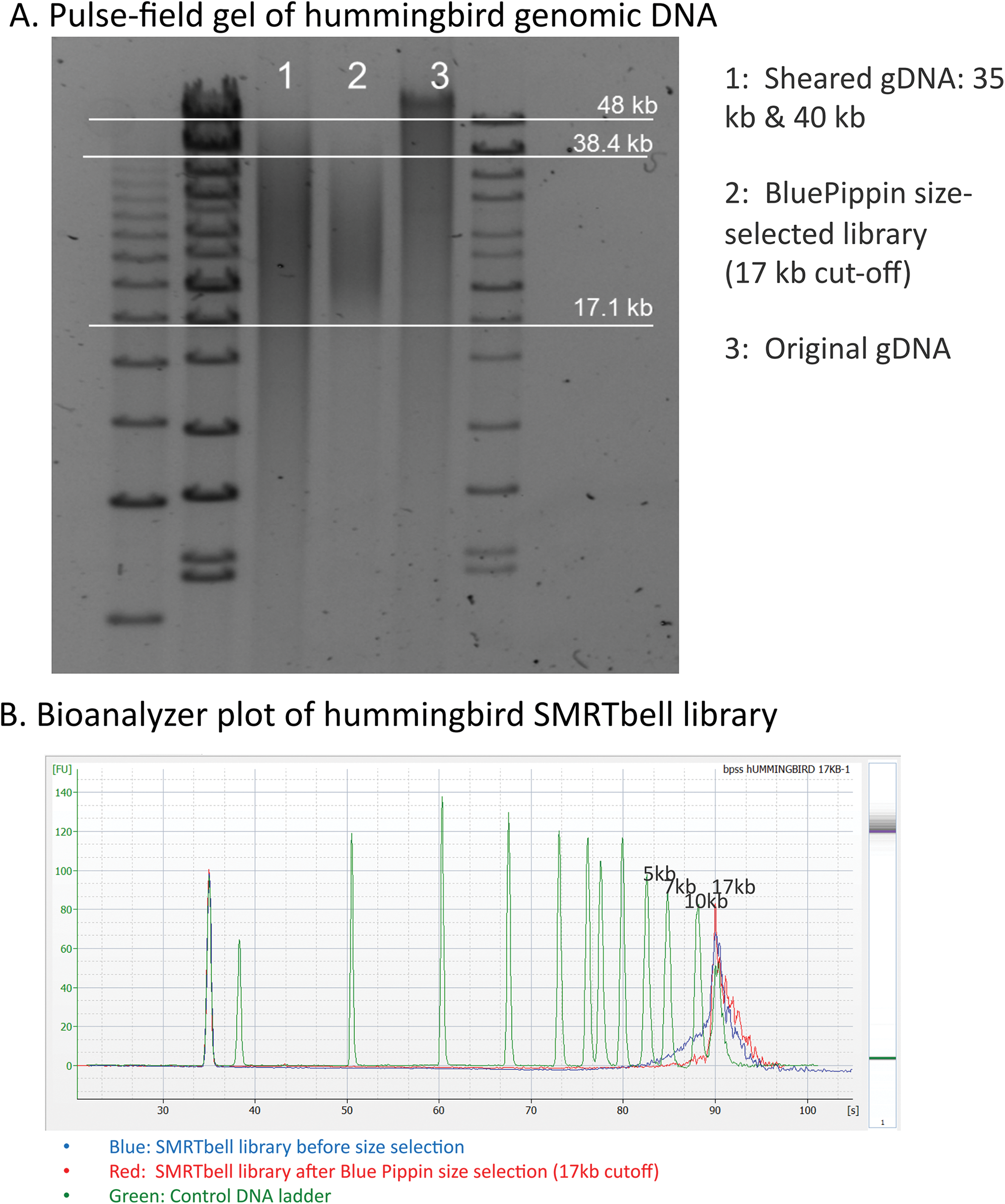
DNA isolation, library construction, and size selection. *(A)* Pulsed-field gel showing original size of starting genomic DNA (lane 3), the sheared DNA (1), and the size selected library (2). *(B)* Bioanalyzer trace before (blue) and after (red) library size selection for fragments > 17 kb.

**Supplementary Figure 2.**
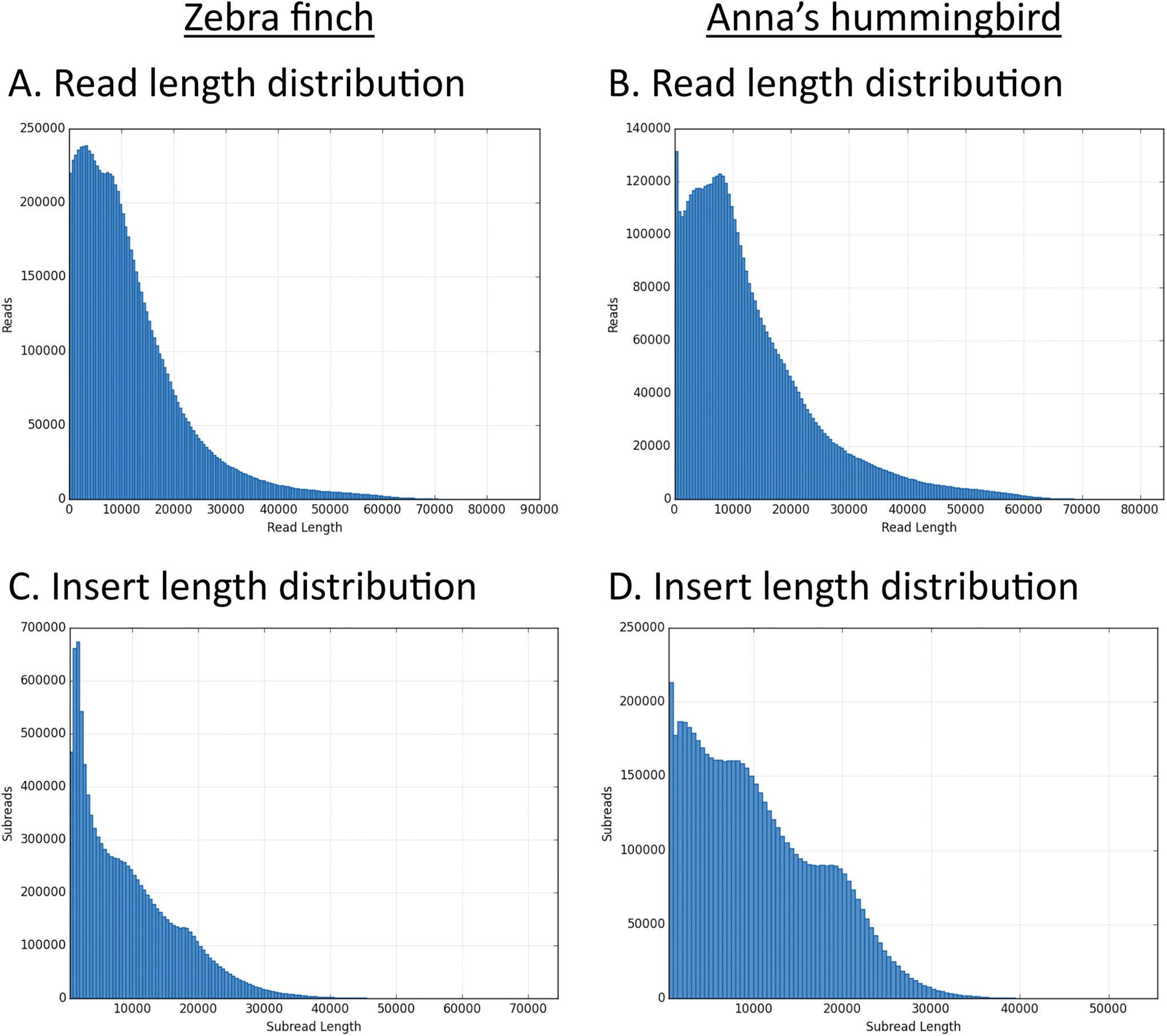
Read and insert length distributions. *(A, B)* Sequence read length distributions from SMRT cell sequencing for both species. *(C, D)* Sequenced DNA insert length distributions from SMRT cell sequencing for both species.

**Supplementary Figure 3.**
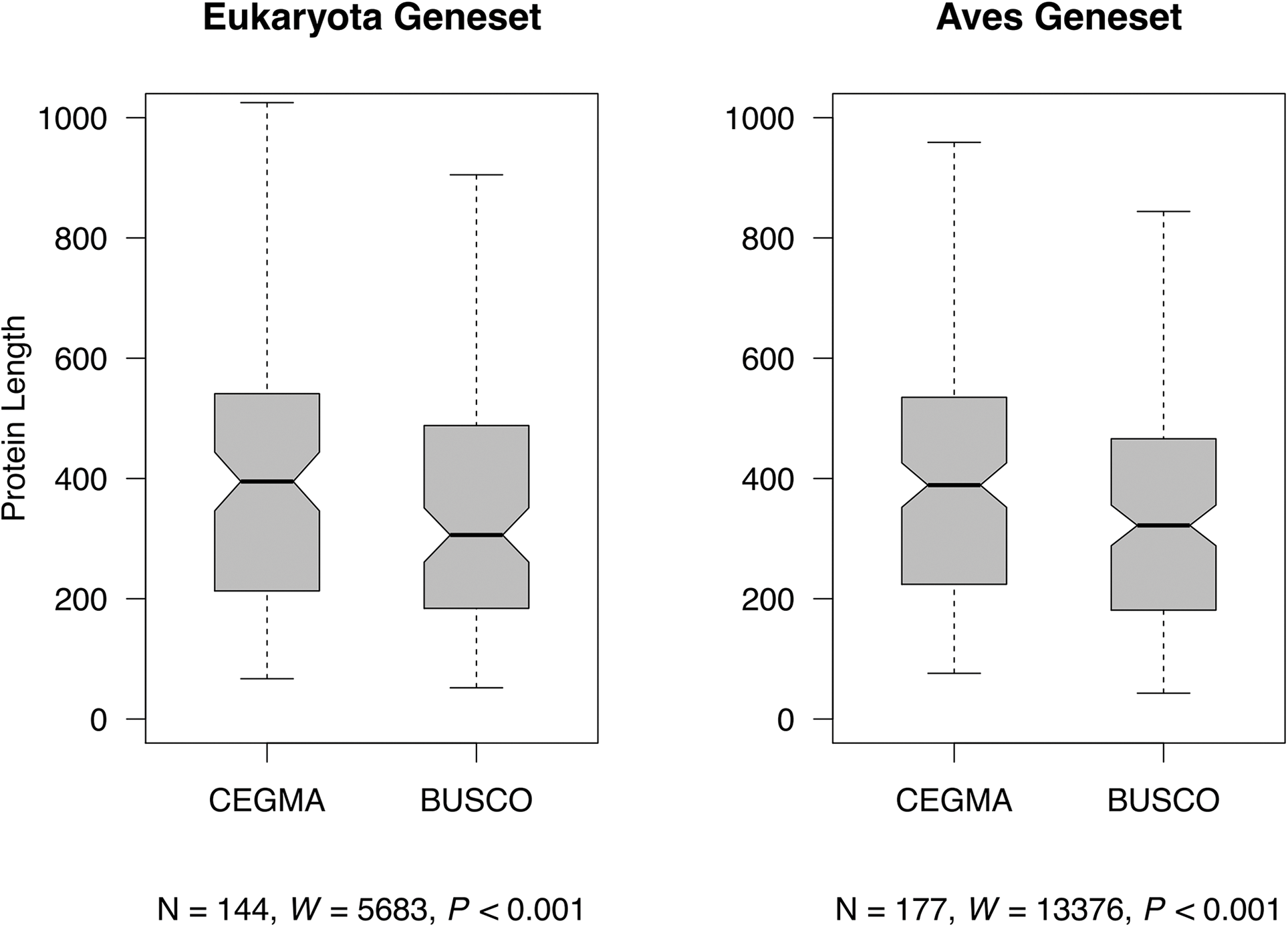
Box plots comparing protein coding sequence lengths of orthologous proteins between the CEGMA and BUSCO eukaryotic and avian datasets. ** p < 0.001; *** p < 0.0001, one-sided paired Wilcoxon signed-rank test, prediction of the proteins being longer in CEGMA datasets.

**Supplementary Figure 4.**
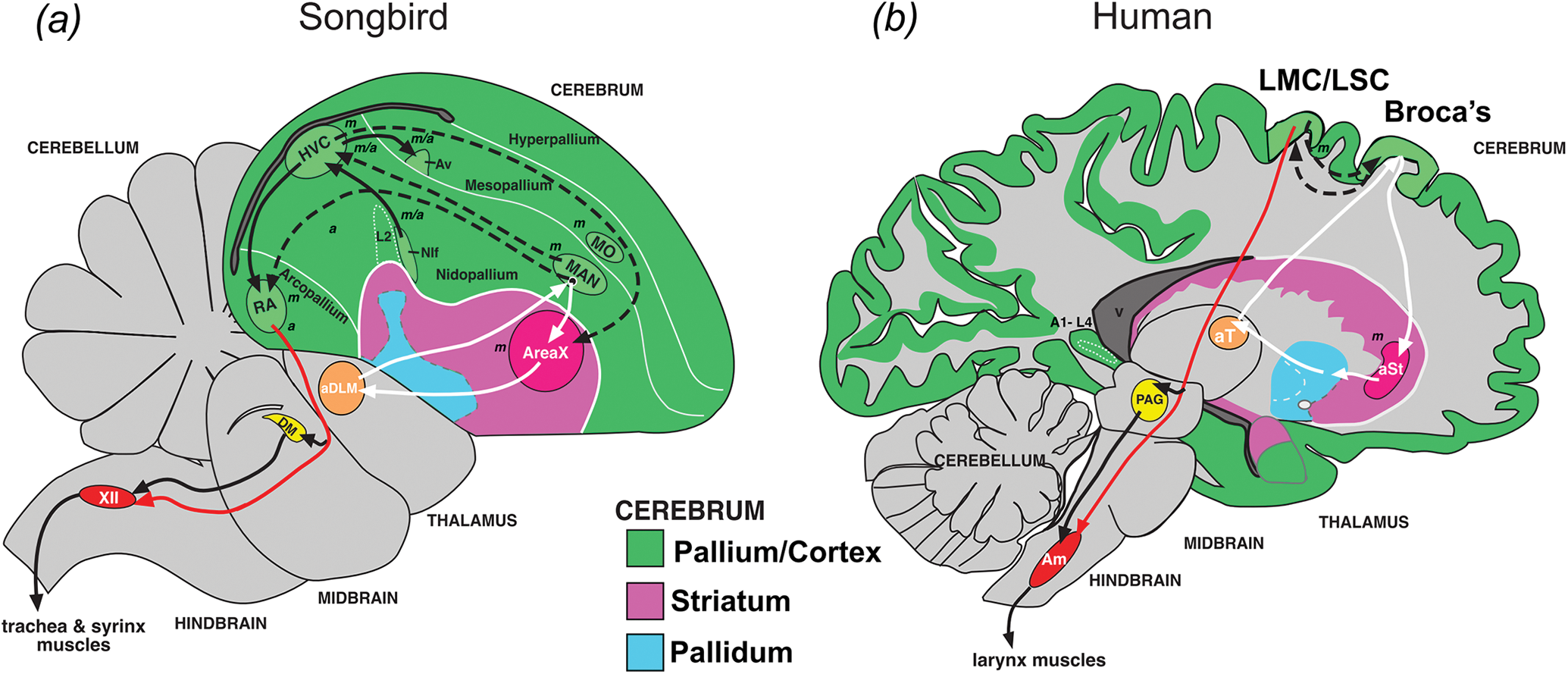
Vocal learning and adjacent brain regions in songbirds used for RNA-Seq and ChIP-Seq analyses, and comparison with humans. *(A)* Drawing of a zebra finch male brain section showing specialized vocal learning pathway and associated profiled song nuclei RA, HVC, LMAN, and Area X. *(B)* Drawing of a human brain section showing spoken-language pathway and analogous brain regions. Black arrows, posterior vocal motor pathway; White arrows, anterior vocal learning pathway; Dashed arrows, connections between the two pathways; Red arrow, specialized direct projection from forebrain to brainstem vocal motor neurons in vocal learners. Italicized letters adjacent to the song and speech regions indicates regions (in songbirds) that show mainly show motor *(m)*, auditory *(a)*, equally both motor and auditory *(m/a)* neural activity or activity-dependent gene expression. Figure from (Chakraborty and Jarvis 2015) and (Pfenning et al. 2014).

**Supplementary Figure 5.**
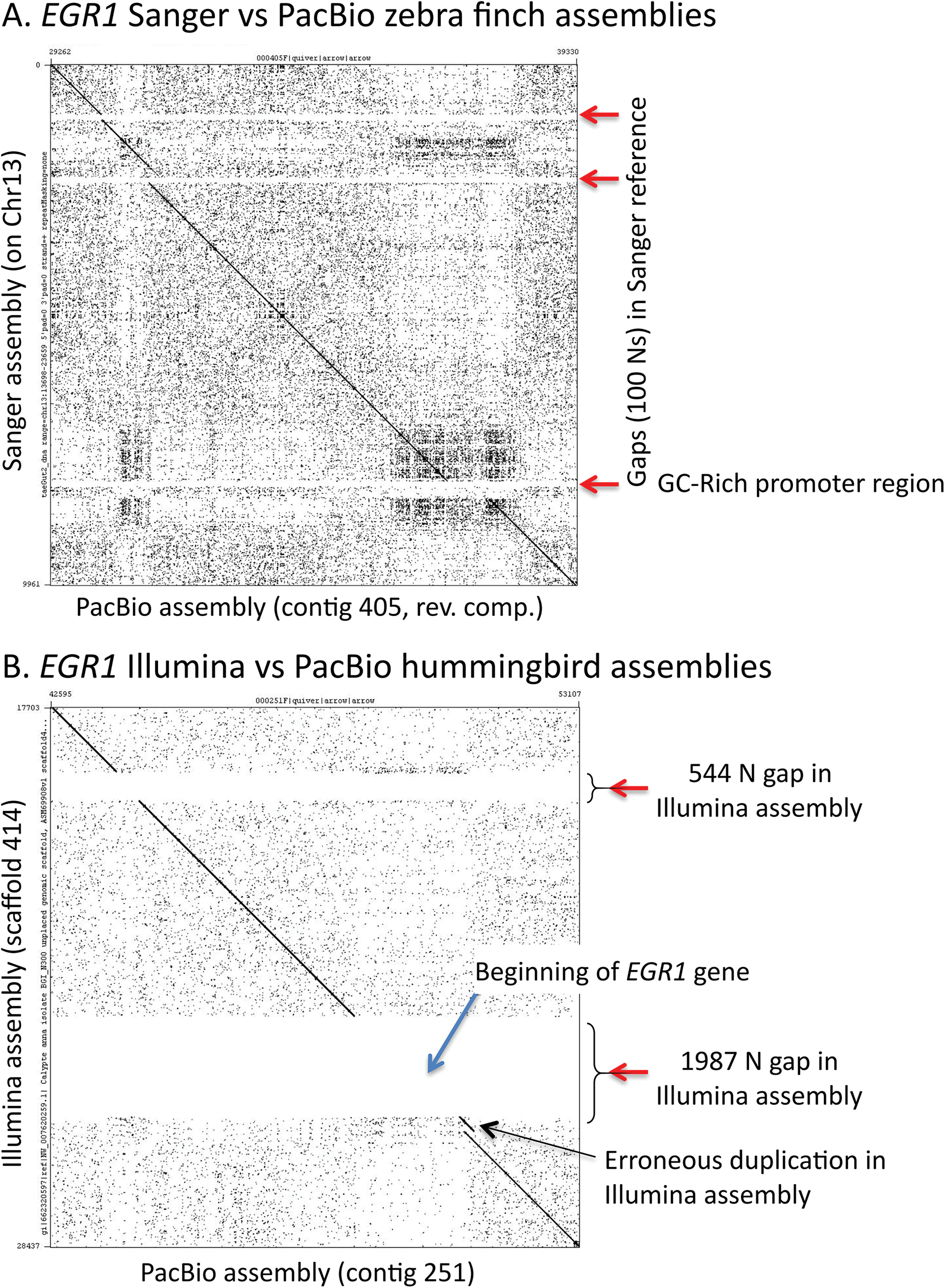
Dot plot of sequence comparisons for genome assemblies of the *EGR1* region. *(A)* Comparison of zebra finch PacBio-based versus Sanger-based assemblies for the region containing *EGR1*, showing the GC-rich promoter region and closing and corrections of gaps for the PacBio-based assembly. *(B)* Comparison of hummingbird Illumina-based versus PacBio-based assemblies for the region containing *EGR1*, showing an erroneous tandem duplication in the Ilumina-based assembly and closing of gaps for the PacBio-based assembly.

**Supplementary Figure 6.**
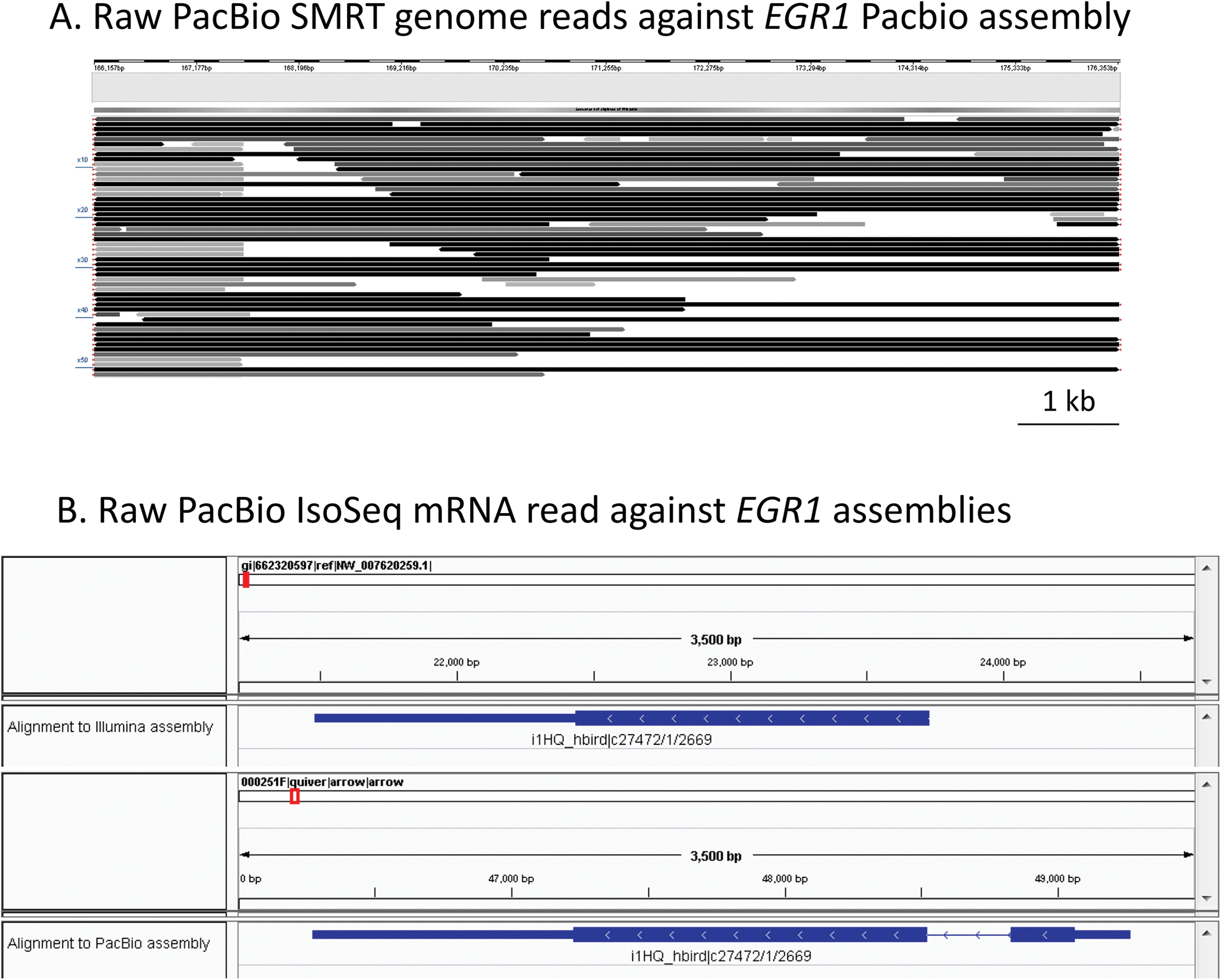
Single SMRT genomic reads and Iso-Seq mRNA reads for *EGR1*. *(A)* Zebra finch PacBio SMRT reads (rows) mapped against the zebra finch PacBio assembly (contig 405, entire *EGR1* region, same as Fig. 3A). Reads are shaded by length (>10 kb reads = black). *(B)* Example of a single Ruby-throated hummingbird Iso-Seq read mapped against Illumina-based (top) and PacBio-based (bottom) Anna’s hummingbird genome assemblies using GMAP. Note the first exon (blue) which is present in the Iso-Seq read is missing in the Illumina-based assembly, but present in the PacBio-based assembly.

**Supplementary Figure 7.**
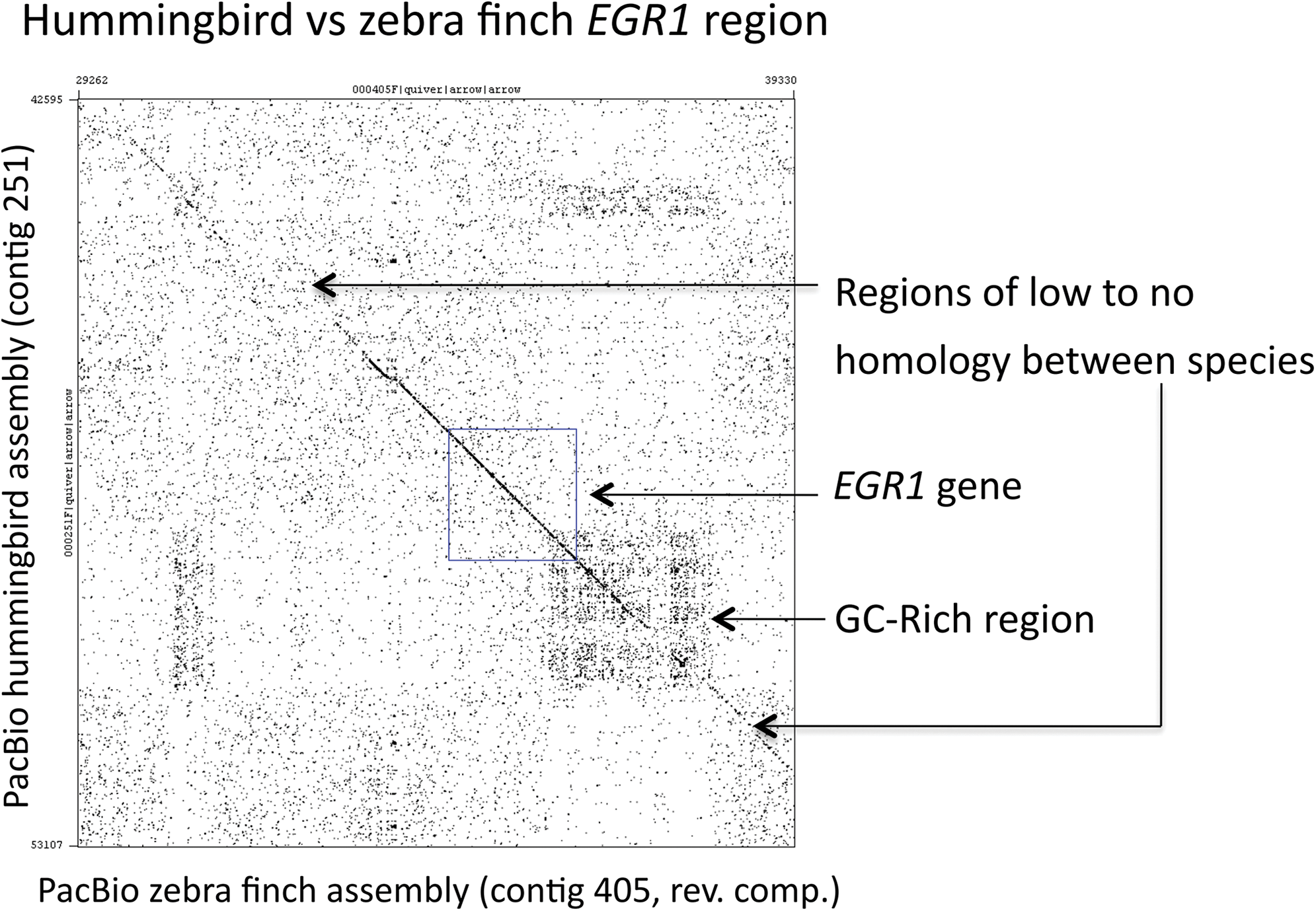
Dot plot of sequence comparison for the PacBio-based hummingbird and zebra finch *EGR1* region assemblies. Note regions of high species conservation and divergence surrounding *EGR1*. Blue box, location of the *EGR1* exons and intron.

**Supplementary Figure 8.**
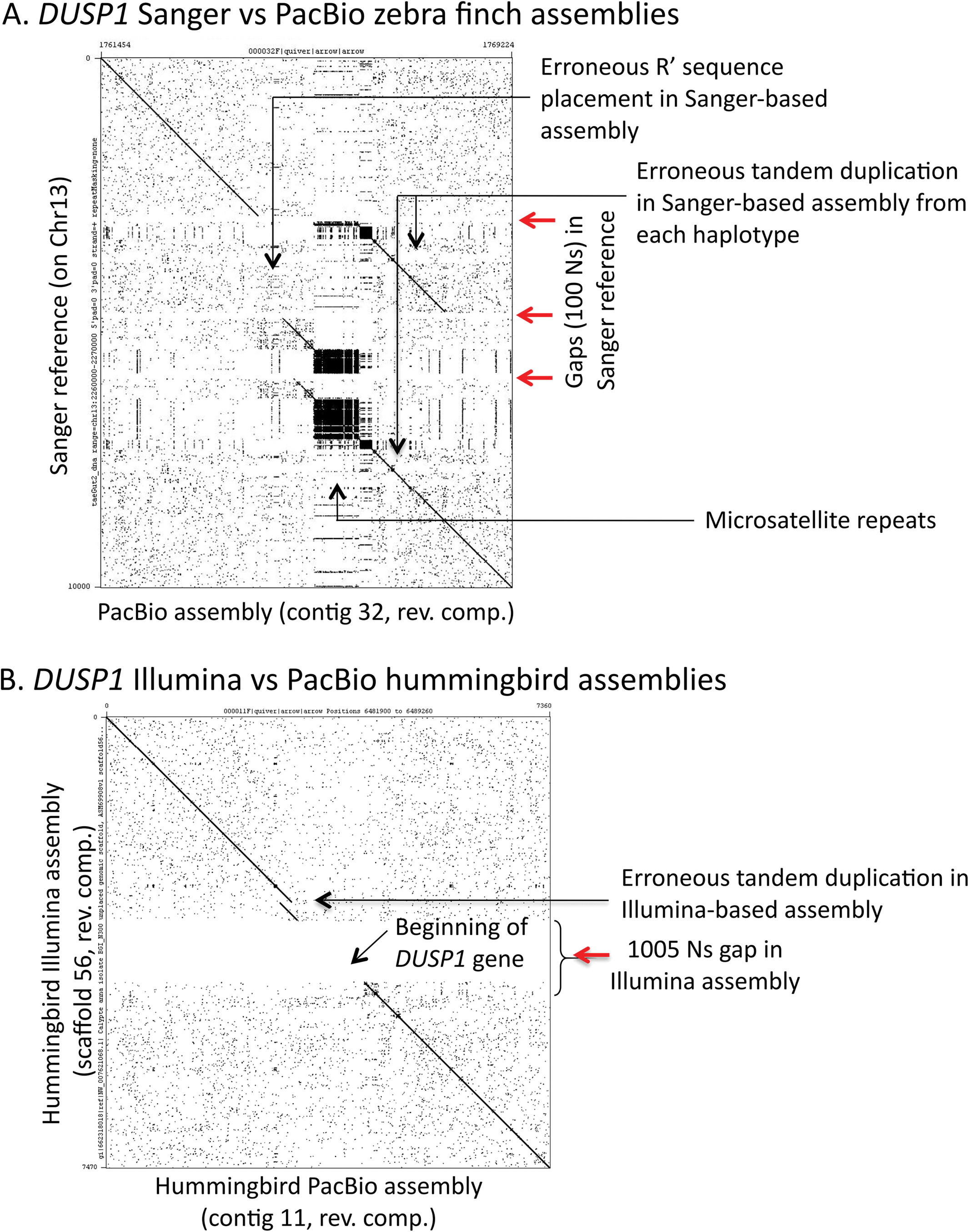
Dot plot comparisons for *DUSP1* region assemblies. *(A)* Comparison of the Sanger-based and PacBio-based zebra finch *DUSP1* region assemblies, showing problems in the Sanger-based assembly with microsatellite repeats. *(B)* Comparison of the Illumina-based and PacBio-based hummingbird *DUSP1* region assemblies, showing a large gap including the microsatellite region and the beginning of the gene, and an erroneous tandem duplication in the Illumina-based assembly.

**Supplementary Figure 9.**
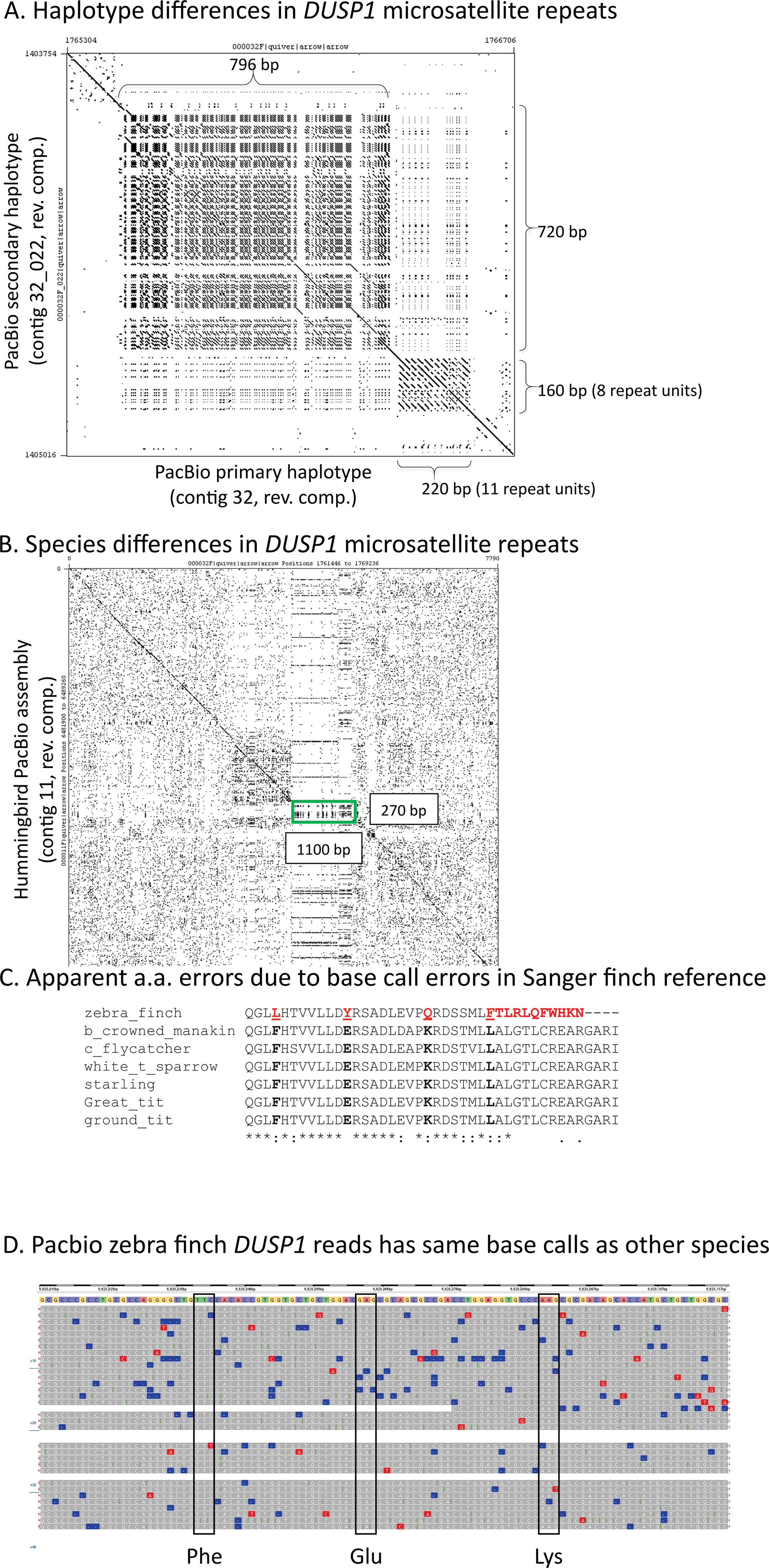
Dot plot comparison of assemblies for the *DUSP1* microsatellite region. *(A)* Differences in the microsatellite region upstream of the *DUSP1* protein coding sequence between the primary and the secondary haplotypes in the fully assembled zebra finch PacBio-based assembly. *(B)* Differences in microsatellites region upstream of *DUSP1* between the zebra finch and hummingbird in the fully assembled PacBio-based assemblies. *(C)* Confirmation of the PacBio sequence in the three locations disparate from the zebra finch reference by alignments to DUSP1 sequences of other songbirds. *(D)* PacBio reads (rows) corresponding to the genomic region in DUSP1 that differs in three location from the zebra finch reference, resulting in a.a. changes. The codons in question are highlighted.

**Supplementary Figure 10.**
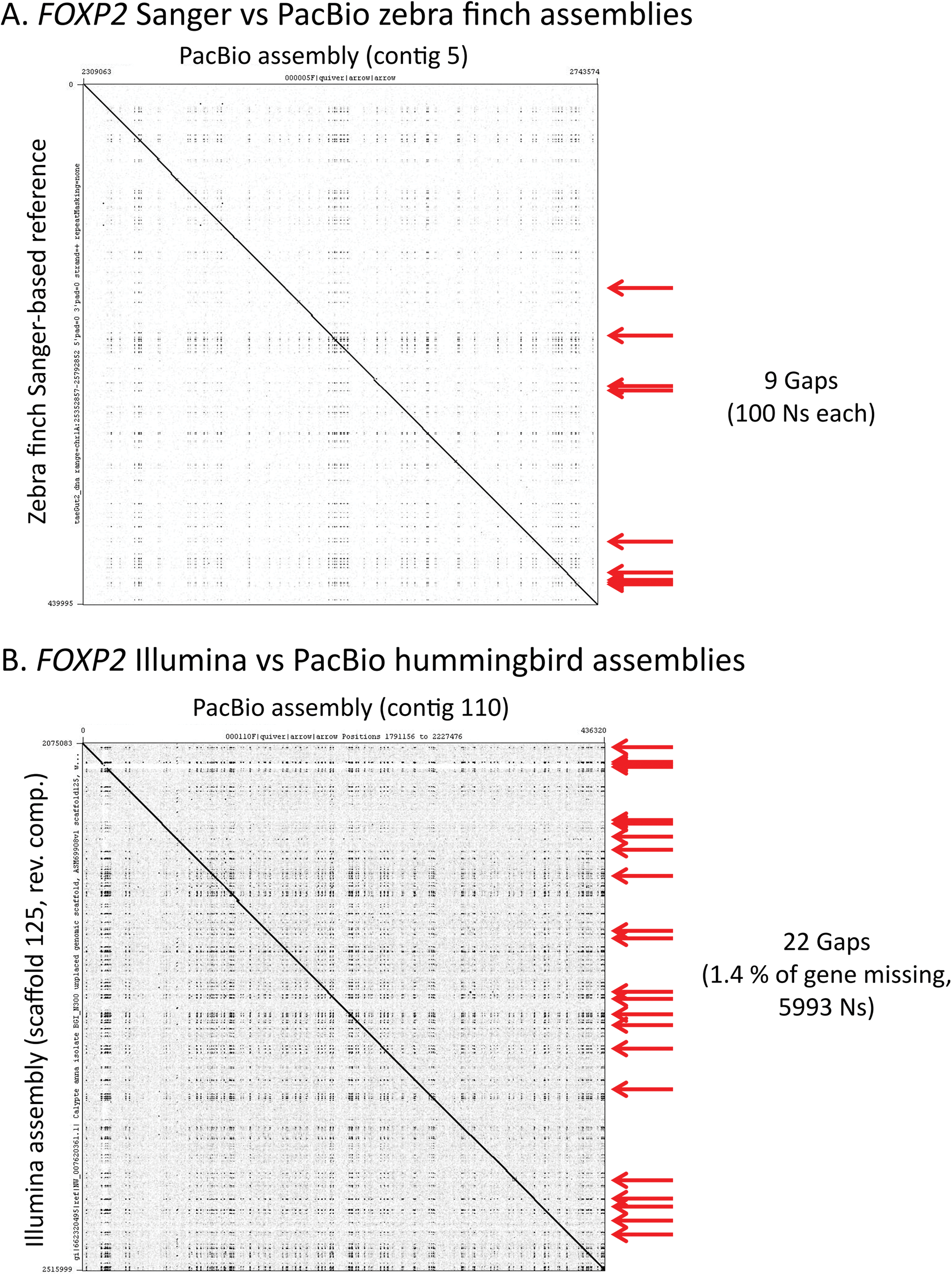
Dot plot comparison of assemblies for the *FOXP2* region. *(A)* zebra finch, *(B)* hummingbird.

**Supplementary Figure 11.**
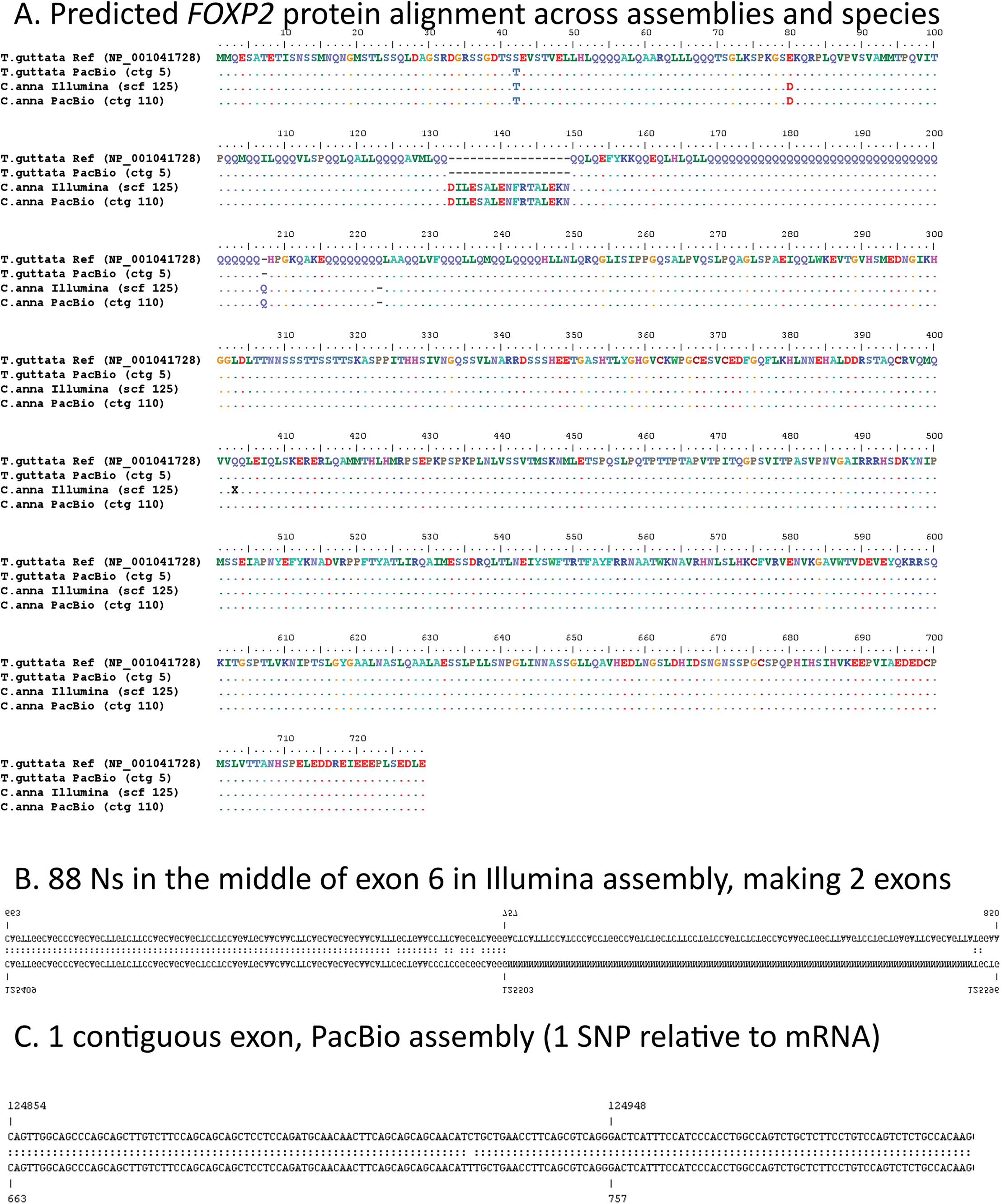
Multiple sequence alignment of the FOXP2 protein for the four assemblies (two zebra finch and two hummingbird) compared in this study, showing correction of a nucleotide error in the Sanger-based zebra finch assembly, and correction of an erroneous stop codon (x) in the Illumina-based hummingbird assembly. Note an extra 18 a.a. stretch in the hummingbird sequence validated by gene prediction of both assemblies, that was not present in the zebra finch.

**Supplementary Figure 12.**
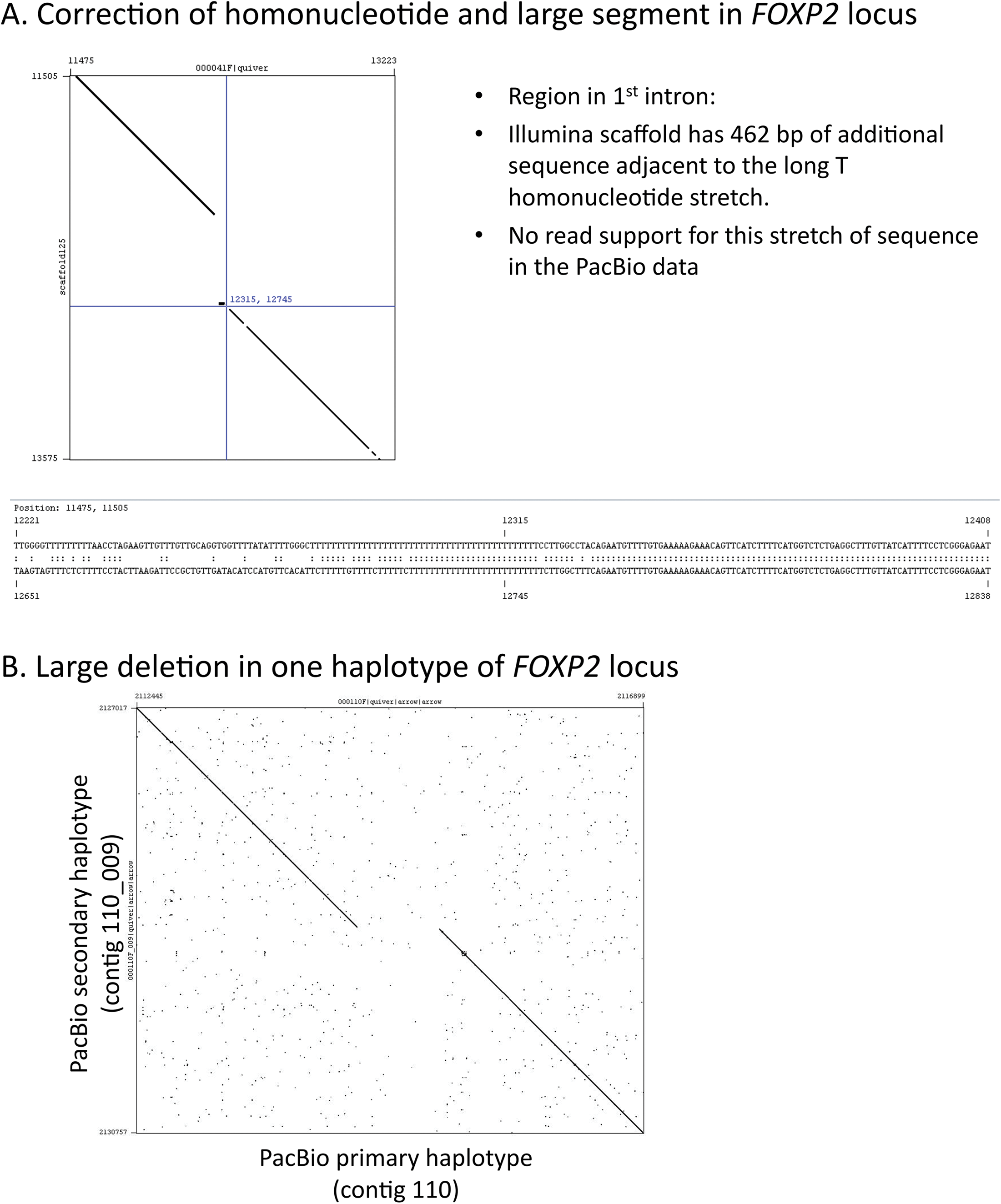
Large regional correction made by the PacBio diploid assembly. *(A)* Correction of an erroneous stretch of 462 bp in the first intron of *FOXP2* in the hummingbird Illumina assembly by the PacBio assembly. *(B)* Dot plot of allelic variation in the *FOXP2* gene by the PacBio diploid assembly: a 708 bp deletion in the secondary haplotype contig relative to the primary contig.

**Supplementary Figure 13.**
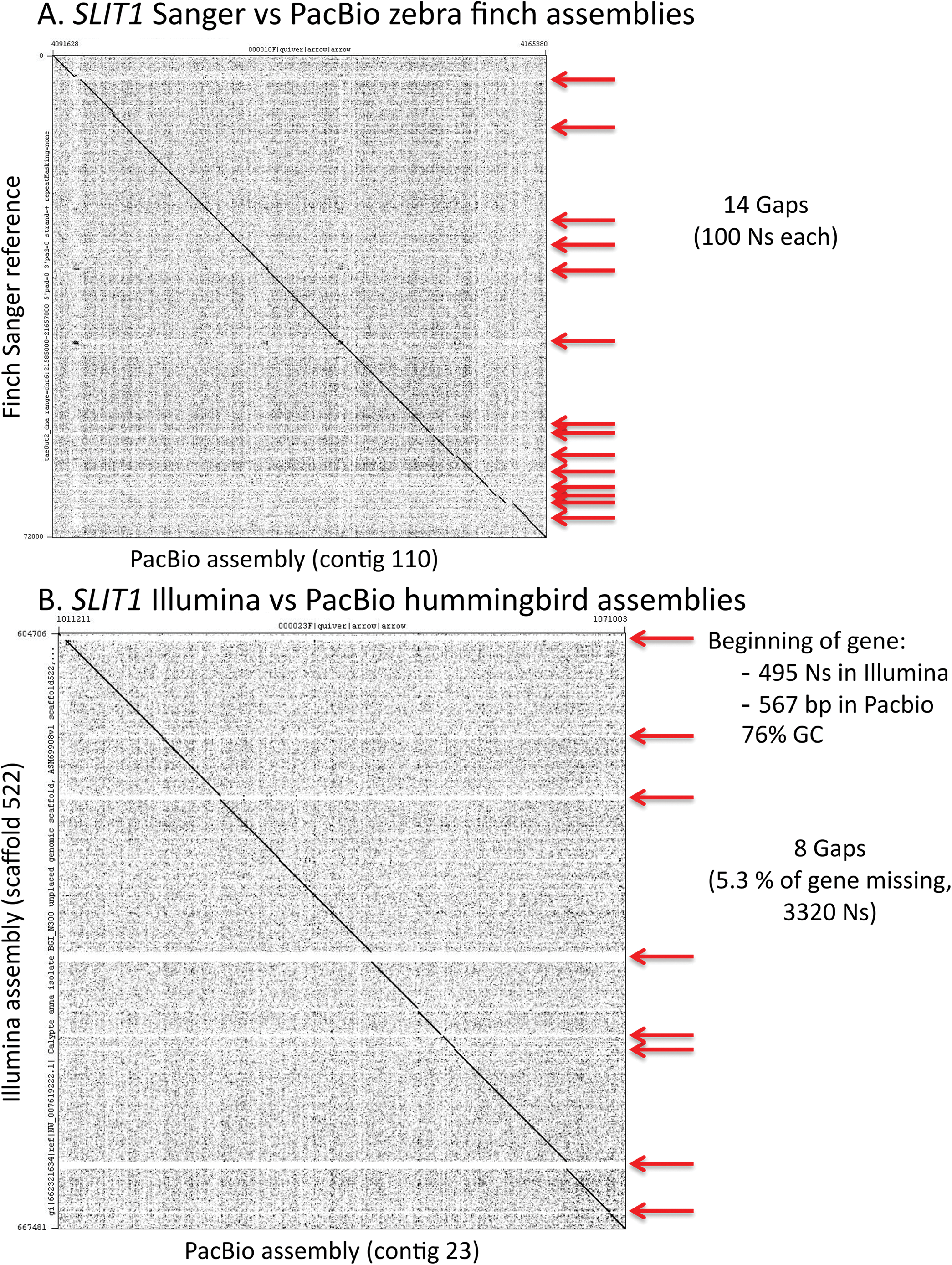
Dot plot comparison of assemblies for the *SLIT1* region. *(A)* zebra finch, *(B)* hummingbird.

